# Host-induced cell wall remodelling impairs opsonophagocytosis of *Staphylococcus aureus* by neutrophils

**DOI:** 10.1101/2023.03.15.532736

**Authors:** Elizabeth V. K. Ledger, Andrew M. Edwards

**Affiliations:** Centre for Bacterial Resistance Biology, Imperial College London, Armstrong Rd, London, SW7 2AZ, UK

**Keywords:** *Staphylococcus aureus*, neutrophil, antibody, peptidoglycan, opsonophagocytosis, immune evasion

## Abstract

The bacterial pathogen *Staphylococcus aureus* adapts to the host environment by increasing the thickness of its cell wall. However, the impact of cell wall thickening on susceptibility to host defences is unclear. Here, we show that as bacteria adapted to serum, the resulting increase in cell wall thickness led to a reduction in the exposure of bound antibody and complement and a corresponding reduction in phagocytosis and killing by neutrophils. The exposure of opsonins bound to protein antigens or LTA were most significantly reduced, whilst opsonisation by IgG against wall teichoic acid or peptidoglycan was largely unaffected. Partial digestion of accumulated cell wall in host adapted cells using the enzyme lysostaphin restored opsonin exposure and promoted phagocytosis and killing. Concordantly, the antibiotic fosfomycin inhibited cell wall remodelling and maintained full susceptibility of *S. aureus* to opsonophagocytic killing by neutrophils. These findings reveal that the adaptation of *S. aureus* to the host environment reduces the ability of the immune system to detect and kill this pathogen through reduced exposure of protein- and LTA-bound opsonins via cell wall remodelling.

## Introduction

*Staphylococcus aureus* frequently infects wounds caused by surgery or insertion of intravenous access devices [1,2]. These infections can result in *S. aureus* seeding into the bloodstream, leading to bacteraemia and subsequent metastatic dissemination to sites including the heart, bones and joints [1,3,4,5]. Despite antibiotic therapy and a potent immune response, *S. aureus* infections have a high rate of relapse and frequently become chronic or recurrent [3,5].

Neutrophils are a key host defence against *S. aureus* infection and are recruited to the site of infection from the bloodstream [6,7,8,9,10,11]. The detection of *S. aureus* by neutrophils is largely dependent upon opsonisation of bacteria by bound antibody and complement, which is enabled in most people by the presence of antibodies against a range of different staphylococcal surface structures, including wall and lipoteichoic acids (WTA, LTA), peptidoglycan, capsular polysaccharide and proteins [10,11,12,13,14,15,16,17,18]. Whilst the precise abundance of antibody against each of the major surface structures varies from person to person, antibody against each macromolecule has been demonstrated to be sufficient to trigger opsonophagocytosis [14,15,16,17,18,19,20].

The binding of neutrophils to opsonins on the surface of *S. aureus* occurs via dedicated receptors and triggers phagocytosis of the pathogen followed by the subsequent exposure of ingested bacteria to a raft of bactericidal products including reactive oxygen species, antimicrobial peptides and proteases [11,12].

To combat the threat posed by neutrophils, *S. aureus* has evolved numerous mechanisms of evading opsonic complement and antibody [12,13,21,22]. For example, *S. aureus* produces two immunoglobulin binding proteins, Spa and Sbi, that reduce antibody-mediated opsonisation, whilst the production of proteins such as SCIN, Efb and CHIPS reduces complement deposition and activation and detection by immune cells [12,13,21,22,23,24,25,26,27,28,29,30,31]. As such, the bacterial cell surface is a critically important determinant in immune detection of *S. aureus* and efforts by the pathogen to evade surveillance and killing by host defences.

The staphylococcal cell envelope is a dynamic structure that responds to host-induced stresses [32,33,34,35]. Consequently, *S. aureus* has a thicker cell wall *in vivo* than when growing *in vitro* [36], a phenotype that is replicated when staphylococci are exposed to human serum or present within endothelial or osteoblast cells [37,38,39,40]. In the case of serum, cell wall thickening is triggered when *S. aureus* detects the presence of the host defence antimicrobial peptide LL-37 via the GraRS two component system [37]. This results in significantly greater quantities of both peptidoglycan and WTA in the cell wall, relative to bacteria grown in laboratory culture medium [37]. Importantly, the changes to the cell envelope triggered by human serum are distinct from those that occur during bacterial entry into stationary phase and are also not triggered by incubation of *S. aureus* in PBS or cell culture medium. i.e. serum-induced changes are not simply due to a lack of nutrients or lack of staphylococcal replication, but represent a specific response to the host environment [37].

Host-induced changes to the cell wall are important for the ability of the pathogen to cause and sustain infection. Cell wall thickening has been shown to reduce susceptibility to antibiotics, whilst mutant strains lacking various cell wall synthetic enzymes are less virulent in infection models [32,34,35,37,40]. However, it is unknown whether host-induced changes to the bacterial cell wall affect the detection and killing of *S. aureus* by the host immune system. To address this, we examined the impact of host-induced changes to the staphylococcal cell envelope on subsequent interactions of *S. aureus* with neutrophils. This revealed that cell wall thickening constitutes a previously unrecognised mechanism of immune evasion that functions by significantly reducing the exposure of opsonins bound to proteins and LTA and thereby reducing opsonophagocytic killing.

## Results

### Host adaptation reduces killing of *S. aureus* by neutrophils

To understand the impact of host adaptation on staphylococcal susceptibility to host defences, we either grew bacteria to exponential phase in tryptic soy broth (TSB) to represent standard laboratory conditions or allowed bacteria to adapt to host conditions using a previously described *ex vivo* human serum model that triggers cell wall thickening [37] (Fig. 1A). This model uses pooled human serum, which avoids variability in anti-staphylococcal antibody levels between donors [14]. Since the serum is not heat inactivated, it contains functional immunoglobulins and complement components.

**Figure 1.**
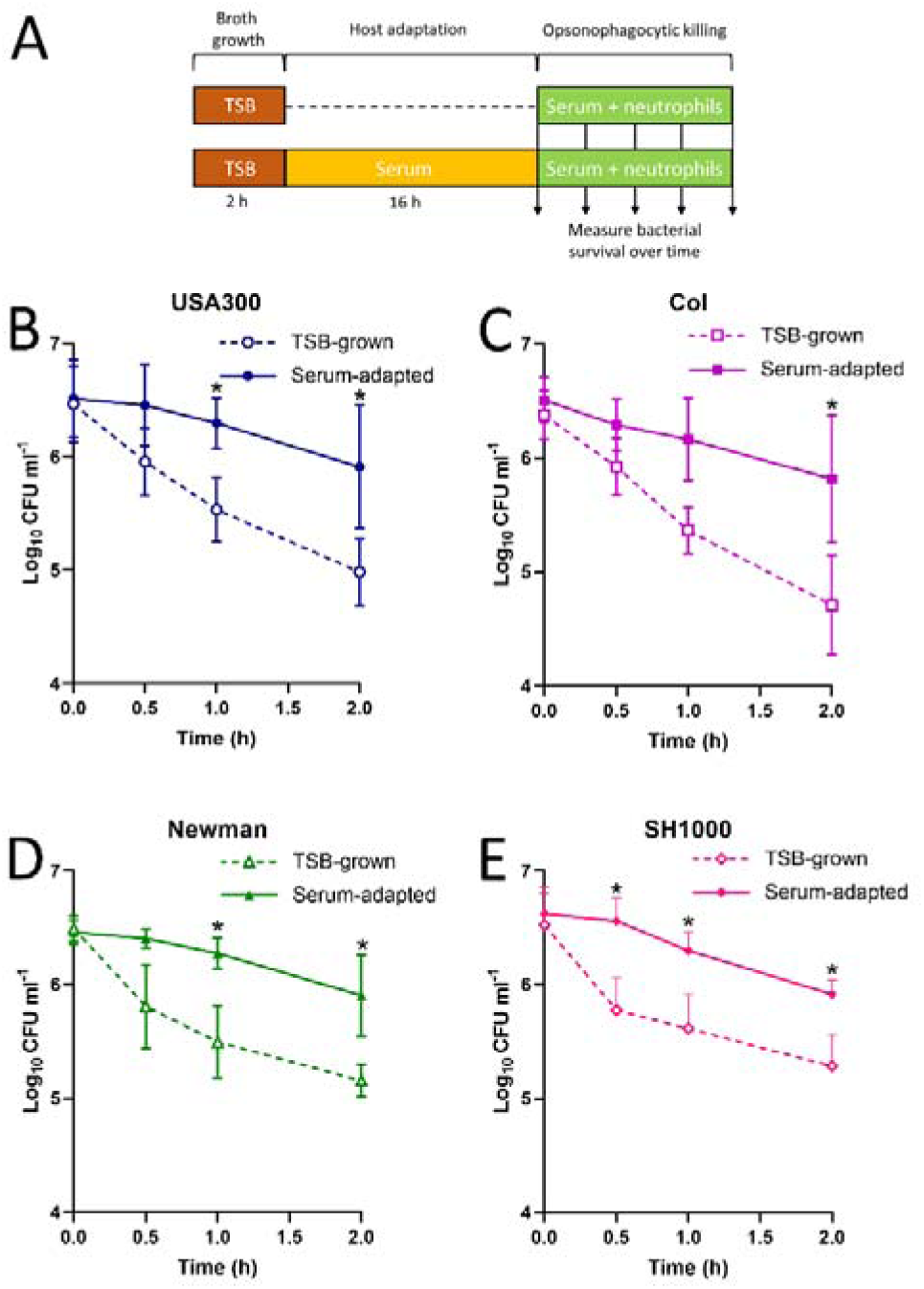
Incubation in human serum promotes tolerance to killing by neutrophils. Bacteria were grown in broth or allowed to adapt to human serum before incubation with human neutrophils in the presence of serum and staphylococcal survival measured over time (**A**). Survival of TSB-grown and serum-adapted cultures of *S. aureus* USA300 (**B**), Col (**C**), Newman (**D**) and SH1000 (**E**) during a 2 h incubation with purified human neutrophils. Data represent the geometric mean ± geometric standard deviation of at least three independent biological replicates. Data were analysed by a two-way ANOVA with Sidak’s *post-hoc* test (*, P < 0.05; Serum-adapted vs TSB-grown at indicated time-points).

In addition to triggering cell wall thickening via the GraRS system, serum also suppresses both the growth of *S. aureus* and activation of the Agr quorum-sensing system that regulates expression of many virulence factors [37,41,42,43,44,45,46,47,48].

We then measured the survival of bacteria prepared under each condition during incubation with purified *ex vivo* human neutrophils from male and female healthy donors in the presence of serum to provide antibody- and complement-mediated opsonisation [50,51]. We examined four distinct wild type *S. aureus* strains to represent both methicillin resistant (USA300, Col) and methicillin susceptible (SH1000, Newman) organisms [52,53,54,55].

For all four of the *S. aureus* strains tested, exponential phase bacteria were efficiently killed over time, with <5% of bacteria remaining viable after 2 hours incubation with neutrophils (Fig. 1B,C,D,E). However, host-adapted bacteria survived at levels up to 5 times greater than that seen for exponential bacteria for all strains (Fig. 1B,C,D,E). In addition to demonstrating that host adaptation reduces staphylococcal susceptibility to host immune defences, the high level of consistency observed across all four strains indicated that this is a conserved phenotype.

### Host adaptation reduces opsonin exposure and opsonophagocytosis

Having found that host adaptation reduced staphylococcal susceptibility to host defences, we next determined the mechanism(s) responsible. Given the consistency in survival data across all four strains examined, we focussed on the USA300 lineage since it is both well characterised and clinically important [52].

We started by assessing whether the increase in survival of host-adapted bacteria was due to impaired phagocytosis, using two distinct assays. Bacteria were grown in broth or host-adapted by incubation in serum, before washing in PBS and then incubation with neutrophils in the presence of fresh serum to enable opsonisation (Fig. 2A). The first phagocytosis assay was a flow cytometry-based approach that determined how many fluorescently-labelled bacteria were associated (or not) with neutrophils [56] (Supplementary figure S1). This revealed that the majority of both broth-grown and host-adapted *S. aureus* were phagocytosed after 30 min incubation with neutrophils. However, whilst <3% of broth-grown bacteria remained unphagocytosed, >20% of host-adapted bacteria were free from neutrophils (Fig. 2B). This finding was replicated in a second phagocytosis assay that measured the viability of free and neutrophil-associated bacteria [57], with >10% of host-adapted *S. aureus* unphagocytosed compared with <1% of broth-grown *S. aureus* cells (Fig. 2C; Supplementary figure S2). Combined, these two assays demonstrated that host-adapted *S. aureus* were significantly better at evading phagocytosis than broth-grown bacteria.

**Figure 2.**
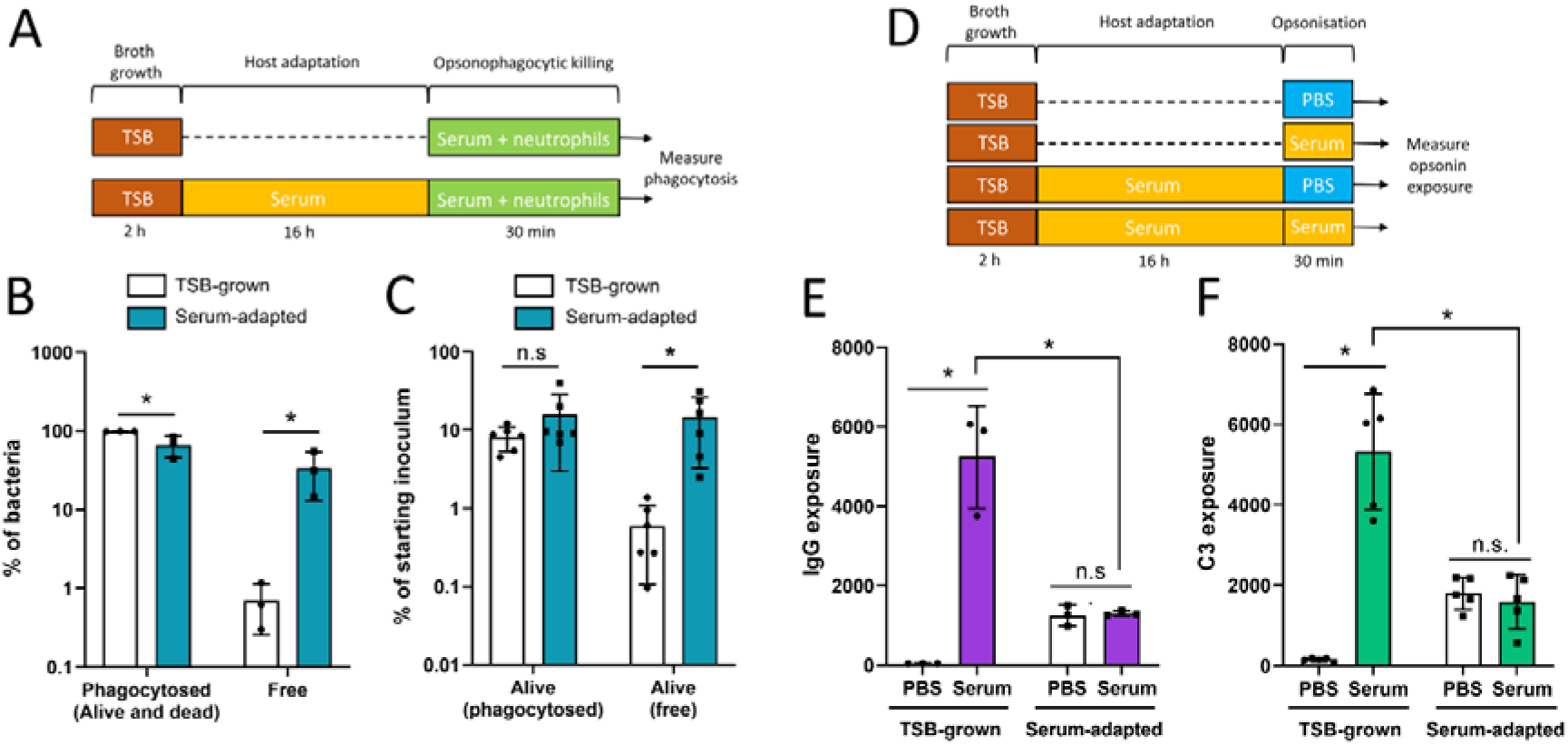
Incubation in human serum reduces opsonophagocytosis and surface-exposed IgG and complement. TSB-grown and serum-adapted cultures of *S. aureus* USA300 were incubated with purified human neutrophils for 30 min (**A**) before the percentage of neutrophil-associated (phagocytosed) and non-neutrophil associated (free) bacteria were determined by (**B**) flow cytometry and (**C**) CFU counts. TSB-grown and serum-adapted *S. aureus* cells were incubated for 30 min with PBS or 10 % serum (**D**) before the levels of surface-exposed bound (**E**) IgG and (**F**) C3 were determined by flow cytometry. Data in **B** and **C** show values for each independent experiment, with bars representing the mean ± standard deviation. In **E** and **F**, each biological repeat is represented by the median fluorescence value of 10,000 bacterial cells. Data in **B, C, E, F** were analysed by two-way ANOVA with Sidak’s *post-hoc* test (*, P < 0.05; n.s, P ≥ 0.05 for the indicated comparisons).

An additional finding was that there were equal numbers of viable host-adapted and broth-grown *S. aureus* cells associated with neutrophils. This indicated that host-adapted *S. aureus* were as susceptible to the microbicides produced by the neutrophils as broth-grown cells (Fig. 2C). Therefore, we concluded that the enhanced survival of host-adapted bacteria compared with broth-grown bacteria (Fig. 1) was due to enhanced evasion of phagocytosis, rather than resistance to the antibacterial products of neutrophils.

To understand why more host-adapted *S. aureus* were able to evade phagocytosis compared with broth-grown cells, we examined the degree of opsonisation of bacteria by antibody and complement using western blotting. In keeping with previous work [58,59], for this experiment we used a mutant strain of USA300 lacking Spa and Sbi to avoid interference caused by these immunoglobulin binding proteins (Supplementary Fig. S3). Broth-grown or host-adapted bacteria were washed in PBS and then incubated, or not, in fresh serum to enable opsonin binding as used in the opsonophagocytosis assays described above (Fig. 1A) before detection of bound antibody and complement component C3 (Fig. 2D).

Despite their reduced phagocytosis by neutrophils, there was more antibody and complement bound to host-adapted cells than to broth-grown, suggesting that a lack of bound opsonins did not explain the immune evasion phenotype of host-adapted bacteria (Supplementary Fig. S4).

To understand why host-adapted cells had high levels of bound antibody and complement but low levels of phagocytosis, bacteria were prepared as described above for opsonophagocytosis assays and then the levels of surface-exposed antibody and the complement component C3 quantified using flow cytometry (Fig. 2D; Supplementary Fig. S5). Broth-grown bacteria that had been exposed to PBS instead of serum acted as a negative control and confirmed that antibodies used in the assay did not bind non-specifically to *S. aureus* cells (Fig. 2D,E,F). We then showed that, as expected, broth-grown bacteria that were incubated in human serum for 30 min were very strongly bound by both IgG and the complement component C3 (Fig. 2E,F).

Next, we examined host-adapted bacteria and found that they had a significantly reduced level of exposed opsonins, compared with TSB-grown cells, regardless of whether they had been incubated in fresh serum for 30 min or not (Fig. 2E,F). Therefore, despite prolonged incubation in serum and high levels of bound antibody and complement (Supplementary Fig. S4), host-adapted cells had significantly reduced exposure of opsonins on their cell surface relative to broth-grown bacteria that had been opsonised.

Taken together, these experiments revealed that host-adapted bacteria are better able to survive exposure to neutrophils than broth-grown *S. aureus* because they are less likely to be phagocytosed, in keeping with the lower surface exposure of bound IgG and complement.

### Cell wall accumulation impairs opsonophagocytosis by concealing IgG bound to LTA and protein

Since the cell envelope of *S. aureus* accumulates peptidoglycan and WTA during incubation in serum [37], we tested whether this concealed some of the bound antibody and complement. To do this, host-adapted bacteria were incubated for 20 min with a range of sub-lethal concentrations of the enzyme lysostaphin, which cleaves peptidoglycan, to partially remove the cell wall. The lysostaphin was then removed by washing and bacterial viability confirmed by CFU counts. This limited cell wall digestion resulted in a significant, dose-dependent increase in exposure of bound IgG and complement, demonstrating that some of the bound opsonins were concealed by the accumulation of cell wall polymers during incubation in serum (Fig. 3A,B).

**Figure 3.**
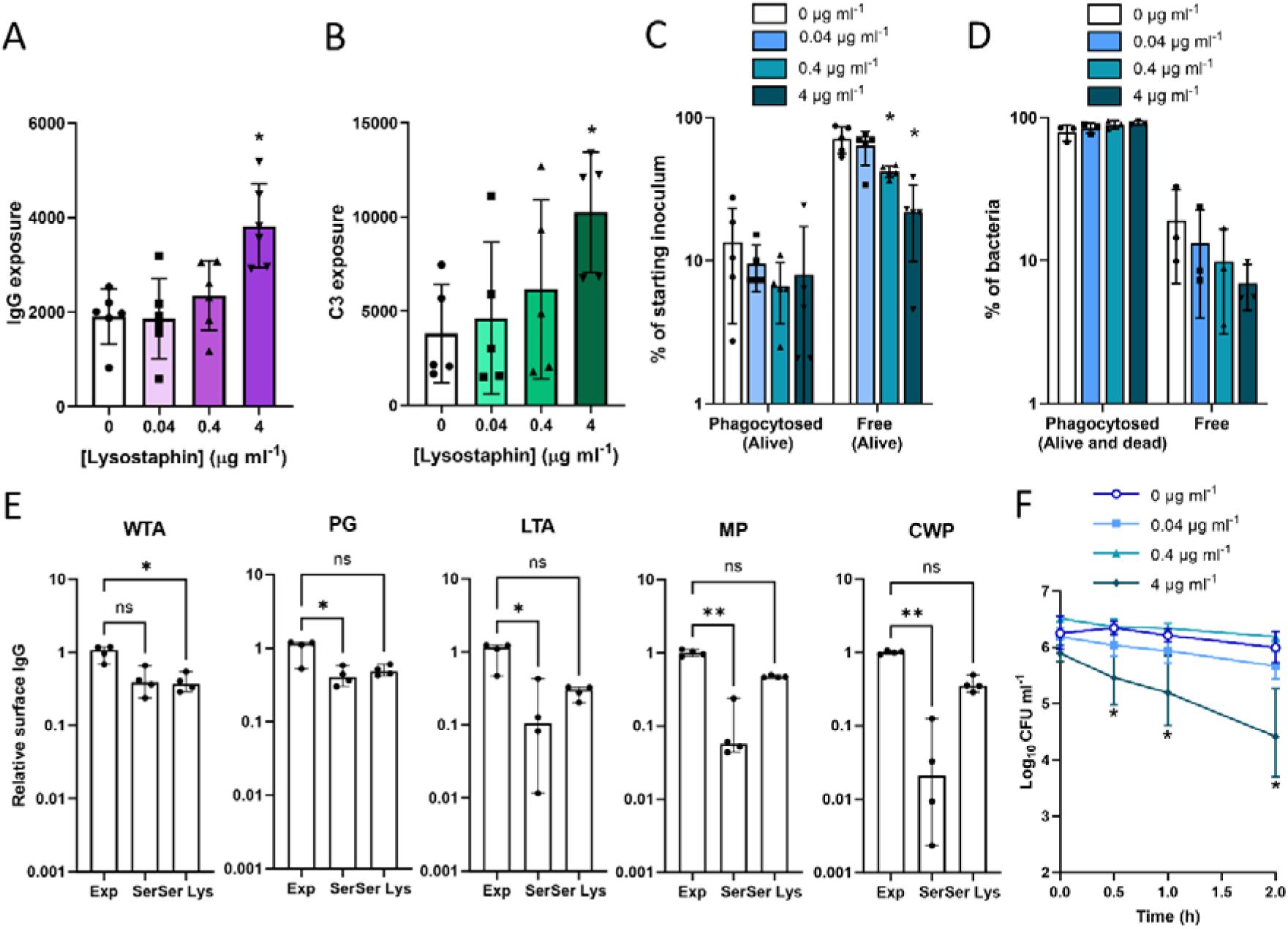
Cell wall accumulation impairs opsonophagocytosis by concealing bound opsonins. Host-adapted cultures of *S. aureus* USA300 were treated with indicated concentrations of lysostaphin for 20 min to partially digest the cell wall before levels of surface-exposed (**A**) IgG and (**B**) complement component C3 were determined by flow cytometry. Additionally, these cells were incubated with purified human neutrophils for 30 min before the percentage of neutrophil-associated (phagocytosed) and non-neutrophil associated (free) bacteria were determined by (**C**) CFU counts and (**D**) flow cytometry. (**E**) Binding ligands of IgG eluted from the surface of *S. aureus* grown to exponential phase and then incubated in serum for 30 min (Exp), allowed to adapt to serum (Ser) or serum-adapted cells that were subsequently subjected to lysostaphin treatment as determined by ELISA. Antibodies were assessed for binding to wall teichoic acid (WTA), peptidoglycan (PG), lipoteichoic acid (LTA), membrane-associated proteins (MP) or cell wall-associated proteins (CWP). (**F**) Survival of lysostaphin-treated cells during a 3 h incubation with purified human neutrophils. Lysostaphin was washed away before incubation with neutrophils. Data in **A** – **D** represent the mean ± standard deviation of the indicated number of independent biological replicates. In **A** and **B**, each biological repeat is represented by the median fluorescence value of 10,000 bacterial events. Data in **A** and **B** were analysed by one-way ANOVA with Dunnett’s *post-hoc* test. Data in **C**, **D** and **F** were analysed by two-way ANOVA with Dunnett’s *post-hoc* test (*, P ≤ 0.05; Lysostaphin-treated vs non-lysostaphin treated). Data in **E** represent the median ± 95% CI of four independent biological replicates and were analysed by Kruskal Wallis test and Dunn’s *post-hoc* test to establish statistically significant differences between groups (**, P < 0.01; *, P < 0.05; ns, P ≥ 0.05 for the indicated comparisons). Data in **F** represent the geometric mean ± geometric standard deviation of 4 independent experiments.

We then tested whether the concealment of bound IgG by accumulated cell wall explained the reduced phagocytosis of host adapted bacteria relative to TSB-grown *S. aureus* . In keeping with increased IgG and complement exposure, limited lysostaphin treatment increased the phagocytosis of host-adapted *S. aureus* by neutrophils (Fig. 3C,D).

Since human serum contains IgG that recognises multiple *S. aureus* surface structures, we next sought to understand whether the reduced opsonisation observed for serum-adapted bacteria was specific to a particular antibody target. Bacteria were grown in TSB and then incubated briefly in serum (30 min) or for 16 h to enable adaptation. Surface exposed IgG was then eluted from bacteria and assessed for its binding to each of the major surface structures by ELISA.

Host-adapted bacteria had similar levels of anti-WTA IgG on their surface compared to exponential phase bacteria and only slightly lower levels of anti-peptidoglycan IgG (2.5-fold difference). However, surface exposure of IgG targeting other surface structures was greatly reduced in host-adapted bacteria compared to exponential phase cells, with anti-LTA IgG 9-fold lower, anti-membrane-associated proteins 18-fold lower and cell wall-associated proteins 48-fold lower (Fig. 3E). We did not examine capsular polysaccharide in these assays as USA300 is deficient in this polymer [52]. As such, the lower surface IgG exposure in host-adapted cells compared to those in exponential phase is primarily due to a loss of exposure of antibody bound to LTA and surface proteins.

Partial digestion of peptidoglycan using lysostaphin restored surface exposure of IgG bound to LTA and proteins to similar levels observed for TSB-grown bacteria (Fig. 3E). Therefore, accumulation of cell wall in host-adapted bacteria preferentially conceals IgG bound to LTA and surface proteins, whilst exposure of anti-WTA and anti-peptidoglycan remains strongly exposed.

Finally, we showed that increasing opsonin exposure via partial lysostaphin digestion of the cell wall, with the enzyme washed away before incubation with immune cells, rendered host adapted staphylococci as susceptible to neutrophil-mediated killing as broth-grown bacteria (Fig. 3F).

Taken together, the experiments described here demonstrate that host-adapted *S. aureus* cells are bound by high levels of antibody and complement but the accumulation of cell wall conceals some of these bound opsonins, reducing phagocytosis and killing by neutrophils.

### Antibiotic-mediated inhibition of peptidoglycan accumulation maintains opsonin exposure and efficient opsonophagocytosis

To further test whether host adaptation reduced phagocytosis via cell wall-mediated concealment of bound opsonins, and to explore potential therapeutic approaches to enhance neutrophil-mediated killing, we first used the antibiotic fosfomycin to block the accumulation of peptidolgycan during host-adaptation, as we have done previously [37]. This antibiotic targets MurA, which catalyses the production of the peptidoglycan precursor UDP *N*-acetylmuramic acid in the cytoplasm [60]. This inhibits peptidoglycan synthesis and prevents serum-induced cell wall thickening from occurring and has been used clinically in anti-staphylococcal combination therapies [37,61,62].

As observed previously, the adaptation of bacteria to serum resulted in a significant reduction in opsonisation, as determined by exposure of IgG and complement, relative to broth-grown bacteria (Fig. 4A,B). However, the presence of fosfomycin in serum significantly reduced opsonin concealment, maintaining IgG and complement exposure at similar levels to that seen for broth-grown bacteria (Fig. 4A,B).

**Figure 4.**
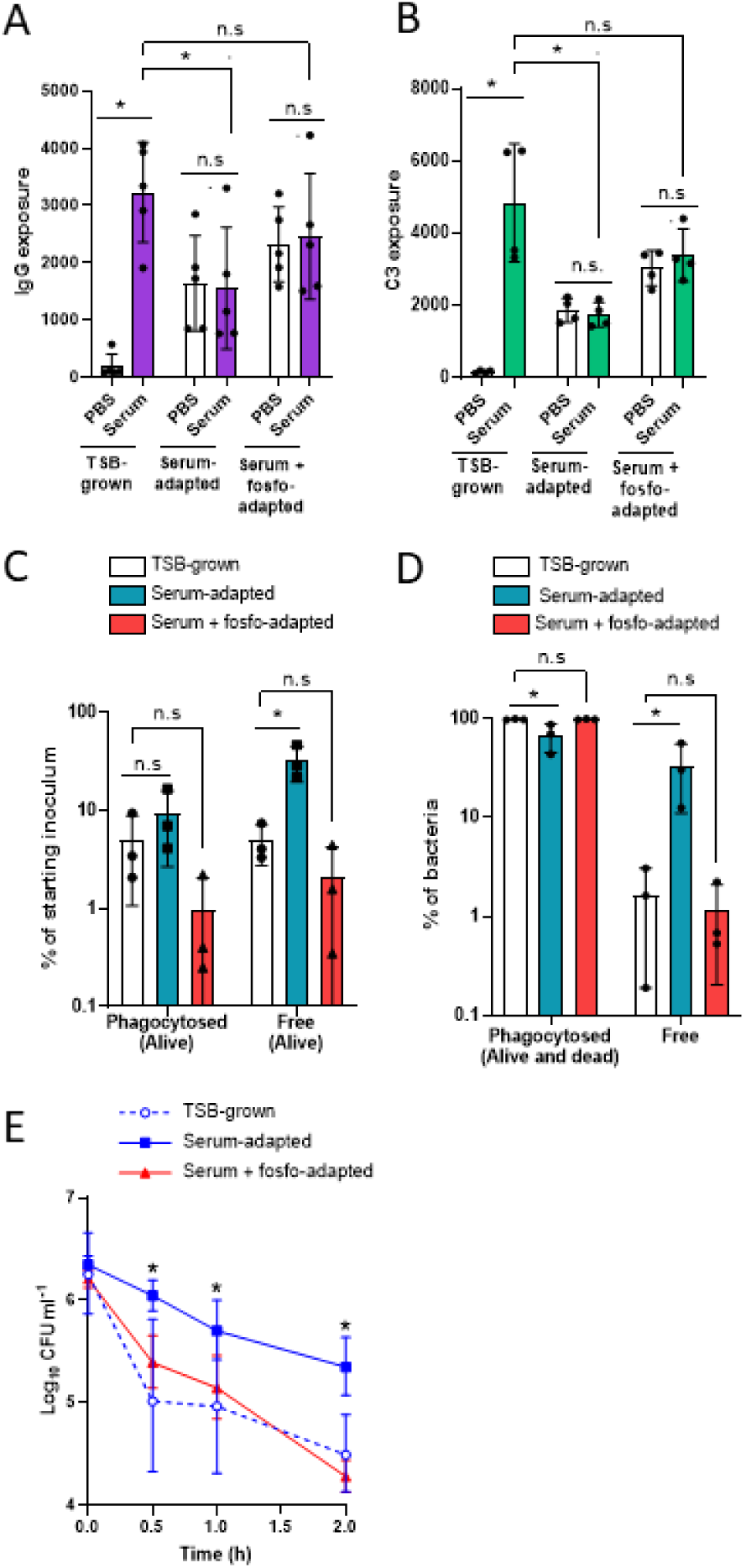
Antibiotic-mediated inhibition of cell wall accumulation maintains opsonin exposure and efficient opsonophagocytosis. TSB-grown, serum-adapted and serum + fosfomycin-adapted cultures of *S. aureus* USA300 were incubated for 30 min with PBS or 10 % serum before the levels of surface-exposed (**A**) IgG and (**B**) complement component C3 were determined by flow cytometry. Additionally, these cells were incubated with purified human neutrophils for 30 min before the percentage of neutrophil-associated (phagocytosed) and non-neutrophil associated (free) bacteria were determined by (**C**) CFU counts and (**D**) flow cytometry. (**E**) Survival of TSB-grown, serum-adapted and serum + fosfomycin-adapted cultures during a 2 h incubation with purified human neutrophils. Summary diagram of the effect of fosfomycin on opsonin exposure during host-adaptation, with antibiotic-mediated inhibition of peptidoglycan accumulation preserving the exposure of bound antibody and complement (**F**). Data in **A** – **D** represent the geometric mean ± geometric standard deviation of the indicated number of independent biological replicates. Data in **E** are presented as the geometric mean ± geometric standard deviation of three independent experiments. In **A** and **B**, each biological repeat is represented by the median fluorescence value of 10,000 bacterial events. Data in **A** and **B** were analysed by two-way ANOVA with Tukey’s *post-hoc* test. Data in **C** – **E** were analysed by two-way ANOVA with Dunnett’s *post-hoc* test (*, P < 0.05; n.s, P ≥ 0.05; comparisons are indicated in **A** – **D** and serum/serum + fosfomycin-adapted vs TSB-grown at each time-point in **E**).

Further analysis of IgG exposure confirmed that fosfomycin treatment preserved the exposure of IgG bound to all major surface structures relative to bacteria that had not been treated with the antibiotic (Supplementary Fig. S6). Similar findings occurred with another inhibitor of peptidoglycan synthesis, oxacillin, which acts on penicillin binding proteins [63], whereas antibiotics that targeted fatty acid biosynthesis (AFN-1252) or DNA gyrase (ciprofloxacin) did not increase IgG exposure relative to host adapted cells that had not been exposed to antibiotics [64,65] (Supplementary Fig. S6). Therefore, in support of our previous findings, we concluded that the accumulation of peptidoglycan during host adaptation significantly reduces the exposure of IgG bound to LTA and surface proteins.

The increased exposure of IgG and complement on the surface of bacteria incubated in serum containing fosfomycin, restored phagocytosis to levels seen with broth-grown bacteria, as determined by both phagocytosis assays (Fig. 4C,D). Furthermore, bacteria that were adapted in serum with fosfomycin were killed by neutrophils as efficiently as broth-grown bacteria, whereas host-adapted bacteria not exposed to fosfomycin survived at significantly higher levels (Fig. 4E). In keeping with our analysis that AFN-1252 did not prevent IgG concealment during host adaptation, this antibiotic did not promote phagocytosis of host-adapted *S. aureus* (Supplementary Fig. S7).

*S. aureus* anchors proteins to peptidoglycan via sortase enzymes (SrtA and SrtB) and so we assessed whether the impact of fosfomycin on host adaption was due to interference with this process. However, host-adapted mutants defective for SrtA or SrtB, which cannot anchor proteins to peptidoglycan, survived incubation with neutrophils as well as host-adapted wild type bacteria (Supplementary Fig. S8A,B).

Taken together, these findings provided additional evidence that cell wall accumulation conceals opsonins bound to LTA and surface proteins, which in turn compromises phagocytosis and killing by neutrophils. They also indicate that the antibiotic fosfomycin, in addition to its own antibacterial activity, may aid the clearance of infection by preventing concealment of opsonins.

## Discussion

The binding of antibody and complement to the bacterial cell surface enables the detection and destruction of pathogens by phagocytic immune cells [10,11,13]. The data presented here demonstrate that *S. aureus* can conceal a subset of bound opsonins via cell wall accumulation, significantly reducing opsonophagocytosis and killing by neutrophils, a previously unrecognised mechanism of immune evasion (Supplementary Fig. S9).

Cell wall remodelling occurs in response to host stresses and provides protection against antibiotics and host defence peptides. It involves the accumulation of peptidoglycan and WTA [36,37,66] and it is therefore unsurprising that exposure of antibody targeting these two polymers was least affected by host adaptation. By contrast, the exposure of antibodies bound to surface proteins and LTA was significantly reduced by host adaptation, in keeping with their localisation within the cell wall itself [67].

Previous work indicated that WTA can block antibodies from binding to antigens within the cell wall [68]. Although WTA accumulates in the wall during host adaptation, it is currently unknown whether this contributes to concealment of IgG bound to LTA or proteins. Unfortunately, since cell wall accumulation is dependent upon D-alanine labelled WTA [37], we could not use a WTA-deficient mutant to explore the role of this polymer in reducing opsonin exposure. However, our work did show that inhibition of peptidoglycan accumulation preserved antibody exposure and opsonophagocytic killing by neutrophils, demonstrating a key role for cell wall accumulation.

During host adaptation, peptidoglycan accumulation is due to a combination of peptidoglycan synthesis and inhibition of autolytic activity [37]. Recent work has revealed that mutants lacking the Atl autolysin have defective surface exposure of staphylococcal surface proteins, inhibiting their recognition by reactive antibodies [69]. Exposure of surface proteins was restored using enzymatic digestion of peptidoglycan, providing additional evidence that peptidoglycan accumulation can obscure surface antigens and prevent their detection by antibodies. However, the impact of concealment of surface proteins on opsonophagocytosis has not been investigated previously.

Several experimental vaccines have been developed based on surface proteins in an attempt to generate high serum titres of opsonising antibody. Unfortunately, despite very promising data from animal infection experiments, none of these vaccines have shown efficacy in humans [70]. Several plausible reasons for this discrepancy have been proposed, including the host-specificity of staphylococcal immune evasion factors and previous staphylococcal infection directing the host towards non-protective immunity [14,70,71].

Another difference between model infection of animals and natural infection in humans is the physiological state of the bacteria. For many animal infections, bacteria are grown in TSB immediately prior to administration into the animal and will therefore have high levels of multiple protein antigens exposed on their surface, which facilitates rapid opsonophagocytosis [72,73,74,74,75]. By contrast, natural invasive infection typically begins with colonisation of superficial sites such as an inserted IV catheter and so bacteria may be host-adapted when they enter the bloodstream and are thus less well recognised by antibodies targeting surface proteins [76,77,78,79,80]. As such, the addition of WTA as a vaccine antigen may provide a reasonable level of protection against bacteria that have accumulated cell wall and thus have reduced exposure of surface proteins.

Previous work has indicated that the thickened cell wall associated with vancomycin resistance reduces staphylococcal susceptibility to intracellular killing by neutrophils [81]. However, our data did not show a difference in staphylococcal survival within neutrophils, with similar numbers of intracellular viable broth-grown and host-adapted bacteria. Instead, the survival advantage of host adaptation appeared to be due to enhanced evasion of phagocytosis.

We do not yet know if these findings are applicable to other Gram-positive pathogens. However, since previous work has shown that serum triggers cell wall thickening in *Enterococcus faecalis* and viridans group streptococci, it is possible that our findings with *S. aureus* represent a broadly conserved mechanism of immune evasion [82].

Cell wall thickening in *S. aureus* is triggered by bacterial sensing of the host defence antimicrobial peptide LL-37 via the GraRS system [37]. Since LL-37 is present in most tissues and amongst the earliest host responses to infection or trauma [81,82,83], we hypothesise that *S. aureus* has evolved to sense this AMP as an early indicator that it is subject to immune attack and provides an opportunity to employ defensive measures against the impending arrival of neutrophils. In support of this hypothesis, GraRS, the two-component system that detects LL-37, is activated in the early stages of staphylococcal skin colonisation, whilst *S. aureus* mutants lacking GraRS are significantly less virulent than wild type strains in invasive infection models [84,85,86].

In addition to providing protection against opsonophagocytosis, LL-37 exposure triggers reduced susceptibility to the antibiotics daptomycin and vancomycin [37,87], suggesting *S. aureus* employs strategies that are broadly protective against the twin threats of host immunity and antibiotic therapy. This is similar to our previous work showing that induction of the *S. aureus* general stress response regulated by the alternative sigma factor SigB can promote survival of bacteria exposed to host defences and various classes of antibiotics [88]. Further support for the link between *S. aureus*-immune interactions and antibiotic tolerance comes from studies showing that oxidative stress conferred by phagocytic cells reduce staphylococcal susceptibility to antibiotics [89,90].

Whilst the immune response may compromise the efficacy of antibiotic therapy under certain circumstances, our study also highlights how antibiotics and the immune response can work synergistically by showing that fosfomycin blocked LL-37 induced cell wall thickening and thereby maintained exposure of bound opsonins, leading to efficient opsonophagocytic killing. Additionally, previous work has suggested that fosfomycin also promotes killing of *S. aureus* via enhanced production of the neutrophil oxidative burst [91]. However, whilst we exposed *S. aureus* to fosfomycin in serum, this was removed by washing prior to incubation with neutrophils and thus does not explain the enhanced killing effect observed in our assays. This strongly suggests that there are at least two mechanisms by which fosfomycin and neutrophils synergise against *S. aureus* and a greater understanding of this may contribute to more effective therapeutic approaches that reduce the high incidence of relapsing or chronic staphylococcal infections [92].

In summary, we show that *S. aureus* cells are heavily opsonised upon initial exposure to serum. However, *S. aureus* responds to serum by accumulating peptidoglycan, which conceals bound opsonins, reducing phagocytosis and killing by neutrophils.

## Materials and methods

### Bacterial strains and growth conditions

Bacterial strains used in this study are shown in Table 1. Strains were grown at 37 °C on tryptic soy agar (TSA) or in tryptic soy broth (TSB) with shaking (180 r.p.m.) supplemented with erythromycin (10 μg ml^-1^) or kanamycin (90 μg ml^-1^) when required.

**Table 1.** Strains used in this study.

| Strain | Relevant characteristics | Reference/source |
| --- | --- | --- |
| USA300 JE2 | LAC strain of the USA300 lineage of community acquired MRSA cured of plasmids | 36 |
| JE2 *sbi::Tn spa::kan* | JE2 strain defective for immunoglobulin-binding proteins Spa and Sbi | This study |
| JE2 *srtA::Tn* | JE2 strain defective for sortase A, which anchors LPxTG motif-containing proteins to peptidoglycan | 93 |
| JE2 *srtB::Tn* | JE2 strain defective for sortase B, which anchors NPQTN motif-containing IsdC protein to peptidoglycan | 93 |
| SH1000 | *rsbU*+ derivative of laboratory strain 8325-4 | 37 |
| Col | Early hospital-associated MRSA isolate, commonly used in laboratory experiments | 38 |
| Newman | Clinical isolate, methicillin susceptible, used widely in animal and laboratory experiments | 39 |

### Construction of strains

The JE2 *sbi*::Tn/*spa*::*kan* double mutant was constructed via transduction of the kanamycin resistance marker from Newman *spa*::*kan* [94] into the *sbi*::Tn mutant present in the NARSA transposon mutant library [93] using ϕ11.

### IgG Fc binding assay

The Fc portion of human Immunoglobulin G (1 mg, Abcam) was labelled with biotin (Thermo Scientific EZ-Link Sulfo-NHS-Biotin) before removal of unbound biotin by dialysis. Labelled Fc portion was then incubated with PBS washed bacterial cells for 30 min (10 µg protein and 10^9^ CFU *S. aureus* in 1 ml PBS). Unbound immunoglobulin fragment was removed by three rounds of washing with PBS before cells were incubated with streptavidin-alkaline phosphatase for 30 min. Cells were then washed with three rounds of PBS before incubation in 200 µl p-Nitrophenol phosphate substrate solution for ELISA (Merck) for 10 min. Cells were then pelleted by centrifugation and the supernatant recovered and A_405_ determined.

### Generation of broth-grown and host-adapted bacterial cultures

To generate broth-grown bacteria, cultures were grown for 16 h in TSB to stationary phase. These were then diluted to 10^7^ CFU ml^-1^ in fresh TSB and incubated for 2 h at 37 °C until 10^8^ CFU ml^-1^ was reached.

To generate host-adapted bacteria, broth-grown cultures were centrifuged (3,200 x *g* for 10 min), resuspended in an equal volume of human serum from human male AB plasma (Sigma) and incubated for 16 h at 37 °C. As *S. aureus* is unable to replicate in human serum, these cultures were also at 10^8^ CFU ml^-1^ [37,47,48,49]. Where appropriate, serum was supplemented with a sub-lethal concentration of fosfomycin (64 μg ml^-1^), oxacillin (128 μg ml^-1^), ciprofloxacin (160 μg ml^-1^) or AFN-1252 (0.15 μg ml^-1^). These concentrations were chosen based on previous work that showed they were the maximum concentration that did not affect staphylococcal viability in serum [37].

Where appropriate, the cell walls of serum-adapted cultures were degraded by lysostaphin. To do this, 1 ml aliquots of serum-adapted bacteria were washed in PBS and resuspended in 1 ml PBS supplemented with indicated concentrations of lysostaphin (between 0.04 and 4 μg ml^-1^). Bacteria were incubated statically for 20 min at 37 °C before being washed by 3 rounds of centrifugation in PBS.

### Purification of neutrophils

Ethical approval for drawing and using human blood was obtained from the Regional Ethics Committee and the Imperial NHS Trust Tissue Bank (REC Wales approval no. 12/WA/0196 and ICHTB HTA license no. 12275). Donors provided informed consent before donating. Neutrophils were extracted from 45 ml human blood, collected in heparin tubes to prevent coagulation. Blood (15 ml) was carefully layered over 30 °C PolymorphPrep (20 ml) and centrifuged for 1 h at 500 x *g* to separate the different cell types. Neutrophils were collected, washed with Hanks balanced salt solution (HBSS) and adjusted to 5 x 10^6^ cells ml^-5^ in HBSS.

### Determination of bacterial killing by neutrophils and phagocytosis by CFU counts

Neutrophils were adjusted to 5 x 10^6^ cells ml^-1^ in HBSS supplemented with 10 % human serum, 0.1 mM CaCl_2_ and 0.1 mM MgCl_2_. In the case of lysostaphin-treated bacteria, 10 % serum was omitted from the HBSS.

TSB-grown/host-adapted bacteria were generated as described above, washed three times in PBS and then added to neutrophils at 5 x 10^6^CFU ml^-1^. Tubes were incubated with end-over-end mixing at 37°C for 3 h and at each time-point (0, 0.5, 1 and 2 h) aliquots were removed, serially diluted 10-fold in PBS and plated to enumerate CFU ml^-1^.

In addition, the number of phagocytosed/unphagocytosed bacteria was also enumerated at the 0.5 h time-point. A 500 μl aliquot of the neutrophil/bacteria mixture was taken and centrifuged at 500 x *g* for 1 min to pellet the neutrophils, along with any neutrophil-associated bacteria. The supernatant (containing unphagocytosed bacteria) was serially diluted 10-fold in PBS and plated for CFU counts and the pellet was resuspended in 500 μl PBS, serially diluted 10-fold in PBS and plated for CFU counts. The CFU ml^-1^ values in the pellet and the supernatant were divided by the CFU ml^-1^ of the starting inoculum to generate the percentage of CFU ml^-1^ phagocytosed and unphagocytosed, respectively.

To validate the experimental conditions used in this assay, two control experiments were run. Firstly, bacteria were prepared as described above were subjected to centrifugation and the CFU counts pre- and post-centrifugation quantified to determine whether bacteria were pulled out of suspension under these conditions. Secondly, to understand whether bacteria associated with neutrophils were intracellular, bacteria were incubated with neutrophils as described above, before subsequent incubation with or without lysostaphin (40 µg ml^-1^) to kill extracellular bacteria. Neutrophils were then washed and CFU counts determined before washing to remove the lytic enzyme. In some experiments neutrophils were lysed with Triton X-100 (0.1%) to determine whether this affected recovery of CFU.

### Measurement of phagocytosis by flow cytometry

To measure phagocytosis by flow cytometry neutrophils and bacteria were prepared as described above except that immediately before bacteria were added to the neutrophils the bacteria were incubated with 10 μg ml^-1^ fluorescein isothiocyanate for 30 min at room temperature and then washed three times in PBS.

As above, 5 x 10^6^ CFU ml^-1^ bacteria were added to 5 x 10^6^ cells ml^-1^ in HBSS supplemented with 10 % human serum, 0.1 mM CaCl_2_ and 0.1 mM MgCl_2_. In the case of lysostaphin-treated bacteria, 10 % serum was omitted from the HBSS. After a 30 min incubation at 37 °C with end-over-end mixing in the dark, cultures were fixed by the addition of an equal volume of 4 % paraformaldehyde (PFA). Samples were then analysed by flow cytometry using an Amnis CellStream. Bacteria were detected using the 488 nm laser and at least 10,000 bacterial events were recorded. Events with FITC ≥ 2 x 10^3^ were counted as bacteria. Events with an FCS of ≥ 3000 were counted as phagocytosed and < 3000 were counted as unphagocytosed/free.

### Measurement of IgG and complement surface exposure by flow cytometry

TSB-grown and serum-adapted cultures were prepared as described above. A *spa/sbi* double mutant was used to prevent non-specific antibody binding. Aliquots (500 μl) were incubated for 30 min at room temperature in either PBS or 10 % human serum. Samples were washed by three rounds of centrifugation in PBS (13,000 x *g* for 1 min) and blocked for 1 h in 4 % BSA in PBS. Samples were washed once in PBS before IgG was detected with a 1:1000 dilution of goat anti-human IgG antibody labelled with the BV421 fluorophore (Jackson ImmunoResearch) or C3 was detected with a 1:1000 dilution of goat anti-human C3 F(ab’)_2_ labelled with FITC (Protos Immunoresearch). Antibody incubations were carried out statically for 1 h at room temperature in the dark. Samples were washed with PBS by three rounds of centrifugation (13,000 x *g* for 1 min) and fixed in 4 % PFA. Samples were analysed by flow cytometry using an Amnis CellStream. IgG was detected using the 405 nm laser and C3 using the 488 nm laser. At least 10,000 bacterial events were recorded and the median value recorded.

### Measurement of IgG and complement by western blotting

Cultures of TSB-grown and serum-adapted bacteria (1 ml at 10^8^ CFU ml^-1^) were prepared as described above, washed by three rounds of centrifugation in PBS (13,000 x *g* for 1 min) and resuspended in 100 μl PBS. A *spa/sbi* double mutant was used to prevent non-specific antibody binding. Lysostaphin (10 μg ml^-1^) was added and bacteria incubated statically for 1 h at 37 °C. Sample buffer (187.5 mM Tris-HCl (pH 6.8), 6 % SDS, 30 % glycerol, 0.03 % bromophenol blue and 15% beta-mercaptoethanol; 50 μl) was added and samples were incubated at 95 °C for 10 min before 15 μl was loaded onto 10 % polyacrylamide gels. Gels were run in Tris-Glycine running buffer (25 mM Tris, 192 mM glycine, 0.1 % SDS, pH 8.4) at 100 V for 10 min followed by 200 V for 50 min before being transferred onto PVDF membranes (10 V for 60 min). Membranes were blocked for 1 h at room temperature in 5 % milk and 1 % BSA in TBST. IgG was detected using 1:10000 dilution of donkey anti-human IgG conjugated to HRP (Abcam) and C3 was detected by 1:5000 dilution of rabbit anti-C3 (Abcam) followed by 1:10000 dilution of goat anti-rabbit IgG conjugated to HRP (Abcam). Blots were developed using SuperSignal West Pico PLUS chemiluminescent substrate (Thermo scientific) and imaged using the Bio-rad ChemiDoc MP imaging system.

### Characterisation of IgG bound to cells

Bacteria were grown to exponential phase and incubated in serum for 30 min or allowed to adapt for 16 h, followed or not by partial cell wall digestion using lysostaphin as described above. Cells (10) were washed three times in PBS before bound antibody was eluted using 200 µl antibody elution buffer (Pierce) for 5 min. Cells were then removed by centrifugation and the eluted antibody solution neutralised with 100 µl protein A binding buffer (Pierce).

To determine the binding ligands of bound antibodies, 10 µg purified cell surface components LTA (Sigma), WTA [37], peptidoglycan [37], membrane proteins [91] or cell wall proteins [91] were immobilised onto the wells of a Nunc Maxisorp ELISA plate by incubation at 4 °C for 16 h. Remaining binding sites were blocked with PBS containing 3 % bovine serum albumin before addition of the eluted antibody samples (200 µl). Wells containing eluted antibodies were incubated at ambient temperature for 1 h, washed three times with PBS and then 200 µl PBS containing anti-human antibodies conjugated to alkaline phosphatase (Abcam, 1:2000 dilution) was added for 1 h. Wells were again washed three times with PBS and bound alkaline phosphatase quantified using a p-Nitrophenol phosphate substrate solution for ELISA (Merck) and A_405_ readings.

### Statistical analyses

CFU counts were log_10_ transformed and displayed as the geometric meanlll±lllgeometric standard deviations [95]. Other data are displayed as the meanlll±lllstandard deviation or median ± 95 % CI. For all experiments, three or more independent replicates were performed as indicated by individual data points. Data were analysed by one-way ANOVA, two-way ANOVA, or Kruskal Wallis, with appropriate post-hoc multiple comparison test as detailed in figure legends using GraphPad Prism (V8.0).

## Acknowledgements

We thank the blood donors, without whom this study would not have been possible. Joan Geoghegan (University of Birmingham) and Angelika Grundling (Imperial College London) are thanked for providing strains. EVKL was supported by a Wellcome Trust PhD Studentship (203812/Z/16/Z). We acknowledge the technical support and the use of equipment from the SAFB Flow Cytometry Facility (Imperial College London). AME acknowledges funding from the Rosetrees Trust and from the Imperial NIHR Biomedical Research Centre, Imperial College London. All authors acknowledge the provision of strains by the Network on Antimicrobial Resistance in *Staphylococcus aureus* (NARSA) Program: under NIAID/ NIH Contract No. HHSN272200700055C.

## Contributions and study design

EVKL and AME designed the experiments, conducted the experiments, analyzed the data, and wrote the manuscript. The funders had no role in the study design, interpretation of the findings or the writing of the manuscript.

## Data availability statement

All data supporting the findings of this study are available within the paper and its Supplementary Information.

## References

1. Tong SY, Davis JS, Eichenberger E, Holland TL, Fowler VG Jr. *Staphylococcus aureus* infections: epidemiology, pathophysiology, clinical manifestations, and management. Clin Microbiol Rev. 28(3):603–61. 2015.

2. Yang Z, Wang J, Wang W, Zhang Y, Han L, Zhang Y, Nie X, Zhan S. Proportions of *Staphylococcus aureus* and methicillin-resistant *Staphylococcus aureus* in patients with surgical site infections in mainland China: a systematic review and meta-analysis. PLoS One. 10(1):e0116079. 2015.

3. Kuehl R, Morata L, Boeing C, Subirana I, Seifert H, Rieg S, Kern WV, Kim HB, Kim ES, Liao CH, Tilley R, Lopez-Cortés LE, Llewelyn MJ, Fowler VG, Thwaites G, Cisneros JM, Scarborough M, Nsutebu E, Gurgui Ferrer M, Pérez JL, Barlow G, Hopkins S, Ternavasio-de la Vega HG, Török ME, Wilson P, Kaasch AJ, Soriano A; International *Staphylococcus aureus* collaboration study group and the ESCMID Study Group for Bloodstream Infections, Endocarditis and Sepsis. Defining persistent *Staphylococcus aureus* bacteraemia: secondary analysis of a prospective cohort study. Lancet Infect Dis. 20(12):1409–1417. 2020.

4. Bergin SP, Holland TL, Fowler VG Jr, Tong SYC. Bacteremia, Sepsis, and Infective Endocarditis Associated with *Staphylococcus aureus* . Curr Top Microbiol Immunol .409:263–296. 2017.

5. Fowler VG Jr, Olsen MK, Corey GR, Woods CW, Cabell CH, Reller LB, Cheng AC, Dudley T, Oddone EZ. Clinical identifiers of complicated *Staphylococcus aureus* bacteremia. Arch Intern Med. 22;163(17):2066–72. 2003.

6. Nguyen TH, Cheung GYC, Rigby KM, Kamenyeva O, Kabat J, Sturdevant DE, Villaruz AE, Liu R, Piewngam P, Porter AR, Firdous S, Chiou J, Park MD, Hunt RL, Almufarriji FMF, Tan VY, Asiamah TK, McCausland JW, Fisher EL, Yeh AJ, Bae JS, Kobayashi SD, Wang JM, Barber DL, DeLeo FR, Otto M. Rapid pathogen-specific recruitment of immune effector cells in the skin by secreted toxins. Nat Microbiol. 7(1):62–72. 2022.

7. DeLeo FR, Diep BA, Otto M. Host defense and pathogenesis in *Staphylococcus aureus* infections. Infect Dis Clin North Am. 23(1):17–34. 2009.

8. Ellson CD, Davidson K, Ferguson GJ, O’Connor R, Stephens LR, Hawkins PT. Neutrophils from p40phox-/- mice exhibit severe defects in NADPH oxidase regulation and oxidant-dependent bacterial killing. J Exp Med. 203(8):1927–37. 2006.

9. Ha KP, Clarke RS, Kim GL, Brittan JL, Rowley JE, Mavridou DAI, Parker D, Clarke TB, Nobbs AH, Edwards AM. Staphylococcal DNA Repair Is Required for Infection. mBio. 11(6):e02288–20. 2020.

10. Scribner DJ, Fahrney D. Neutrophil receptors for IgG and complement: their roles in the attachment and ingestion phases of phagocytosis. J Immunol. 116(4):892–7. 1976.

11. Wright AE, Douglas SR. An experimental investigation of the rôle of the blood fluids in connection with phagocytosis. Proc R Soc Lond. 72357–370. 1904.

12. Leuzzi R, Bodini M, Thomsen IP, Soldaini E, Bartolini E, Muzzi A, Clemente B, Galletti B, Manetti AGO, Giovani C, Censini S, Budroni S, Spensieri F, Borgogni E, Rossi Paccani S, Margarit I, Bagnoli F, Giudice GD, Creech CB. Dissecting the Human Response to *Staphylococcus aureus* Systemic Infections. Front Immunol. 12:749432. 2021.

13. Guerra FE, Borgogna TR, Patel DM, Sward EW, Voyich JM. Epic Immune Battles of History: Neutrophils vs. *Staphylococcus aureus*. Front Cell Infect Microbiol. 7:286. 2017.

14. Holtfreter S, Kolata J, Bröker BM. Towards the immune proteome of *Staphylococcus aureus* - The anti-*S. aureus* antibody response. Int J Med Microbiol. 300(2-3):176–92. 2010.

15. Dryla A, Prustomersky S, Gelbmann D, Hanner M, Bettinger E, Kocsis B, Kustos T, Henics T, Meinke A, Nagy E. Comparison of antibody repertoires against *Staphylococcus aureus* in healthy individuals and in acutely infected patients. Clin Diagn Lab Immunol . 12(3):387–98. 2005.

16. Wergeland HI, Haaheim LR, Natås OB, Wesenberg F, Oeding P. Antibodies to staphylococcal peptidoglycan and its peptide epitopes, teichoic acid, and lipoteichoic acid in sera from blood donors and patients with staphylococcal infections. J Clin Microbiol . 27(6):1286–91. 1989.

17. Kim MJ, Rah SY, An JH, Kurokawa K, Kim UH, Lee BL. Human anti-peptidoglycan-IgG-mediated opsonophagocytosis is controlled by calcium mobilization in phorbol myristate acetate-treated U937 cells. BMB Rep. 48(1):36–41. 2015.

18. Jung DJ, An JH, Kurokawa K, Jung YC, Kim MJ, Aoyagi Y, Matsushita M, Takahashi S, Lee HS, Takahashi K, Lee BL. Specific serum Ig recognizing staphylococcal wall teichoic acid induces complement-mediated opsonophagocytosis against *Staphylococcus aureus* . J Immunol . 189(10):4951–9. 2012.

19. Lee JH, Kim NH, Winstel V, Kurokawa K, Larsen J, An JH, Khan A, Seong MY, Lee MJ, Andersen PS, Peschel A, Lee BL. Surface Glycopolymers Are Crucial for In Vitro Anti-Wall Teichoic Acid IgG-Mediated Complement Activation and Opsonophagocytosis of *Staphylococcus aureus*. Infect Immun. 83(11):4247–55. 2015.

20. Theilacker C, Kropec A, Hammer F, Sava I, Wobser D, Sakinc T, Codée JD, Hogendorf WF, van der Marel GA, Huebner J. Protection against *Staphylococcus aureus* by antibody to the polyglycerolphosphate backbone of heterologous lipoteichoic acid. J Infect Dis. 205(7):1076–85. 2012.

21. de Jong NWM, van Kessel KPM, van Strijp JAG. Immune Evasion by *Staphylococcus aureus*. Microbiol Spectr. 7(2). 2019.

22. Thammavongsa V, Kim HK, Missiakas D, Schneewind O. Staphylococcal manipulation of host immune responses. Nat Rev Microbiol. 13(9):529–43. 2015.

23. Cruz AR, Boer MAD, Strasser J, Zwarthoff SA, Beurskens FJ, de Haas CJC, Aerts PC, Wang G, de Jong RN, Bagnoli F, van Strijp JAG, van Kessel KPM, Schuurman J, Preiner J, Heck AJR, Rooijakkers SHM. Staphylococcal protein A inhibits complement activation by interfering with IgG hexamer formation. Proc Natl Acad Sci U S A. 118(7):e2016772118. 2021.

24. Falugi F, Kim HK, Missiakas DM, Schneewind O. Role of protein A in the evasion of host adaptive immune responses by *Staphylococcus aureus*. mBio. 4(5):e00575–13. 2013.

25. Zhang L, Jacobsson K, Vasi J, Lindberg M, Frykberg L. A second IgG-binding protein in *Staphylococcus aureus*. Microbiology. 144(4):985–991. 1998.

26. Smith EJ, Visai L, Kerrigan SW, Speziale P, Foster TJ. The Sbi protein is a multifunctional immune evasion factor of *Staphylococcus aureus*. Infect Immun. 79(9):3801–9. 2011.

27. de Haas CJ, Veldkamp KE, Peschel A, Weerkamp F, Van Wamel WJ, Heezius EC, Poppelier MJ, Van Kessel KP, van Strijp JA. Chemotaxis inhibitory protein of Staphylococcus aureus, a bacterial antiinflammatory agent. J Exp Med. 199(5):687–95. 2004.

28. Ko YP, Kuipers A, Freitag CM, Jongerius I, Medina E, van Rooijen WJ, Spaan AN, van Kessel KP, Höök M, Rooijakkers SH. Phagocytosis escape by a *Staphylococcus aureus* protein that connects complement and coagulation proteins at the bacterial surface. PLoS Pathog . 9(12):e1003816. 2013.

29. Rooijakkers SH, Ruyken M, Roos A, Daha MR, Presanis JS, Sim RB, van Wamel WJ, van Kessel KP, van Strijp JA. Immune evasion by a staphylococcal complement inhibitor that acts on C3 convertases. Nat Immunol. 6(9):920–7. 2005.

30. Rooijakkers SH, Ruyken M, van Roon J, van Kessel KP, van Strijp JA, van Wamel WJ. Early expression of SCIN and CHIPS drives instant immune evasion by *Staphylococcus aureus*. Cell Microbiol. 8(8):1282–93. 2006.

31. Howden BP, Giulieri SG, Wong Fok Lung T, Baines SL, Sharkey LK, Lee JYH, Hachani A, Monk IR, Stinear TP. *Staphylococcus aureus* host interactions and adaptation. Nat Rev Microbiol. 1–16. 2023.

32. Pidgeon SE, Pires MM. Cell Wall Remodeling of *Staphylococcus aureus* in Live *Caenorhabditis elegans*. Bioconjug Chem. 28(9):2310–2315. 2017.

33. Sobral R, Tomasz A. The Staphylococcal Cell Wall. Microbiol Spectr. 7(4). 2019.

34. Salamaga B, Kong L, Pasquina-Lemonche L, Lafage L, von Und Zur Muhlen M, Gibson JF, Grybchuk D, Tooke AK, Panchal V, Culp EJ, Tatham E, O’Kane ME, Catley TE, Renshaw SA, Wright GD, Plevka P, Bullough PA, Han A, Hobbs JK, Foster SJ. Demonstration of the role of cell wall homeostasis in *Staphylococcus aureus* growth and the action of bactericidal antibiotics. Proc Natl Acad Sci U S A. 118(44):e2106022118. 2021.

35. Monteiro JM, Fernandes PB, Vaz F, Pereira AR, Tavares AC, Ferreira MT, Pereira PM, Veiga H, Kuru E, VanNieuwenhze MS, Brun YV, Filipe SR, Pinho MG. Cell shape dynamics during the staphylococcal cell cycle. Nat Commun. 6:8055. 2015.

36. Sutton JAF, Carnell OT, Lafage L, Gray J, Biboy J, Gibson JF, Pollitt EJG, Tazoll SC, Turnbull W, Hajdamowicz NH, Salamaga B, Pidwill GR, Condliffe AM, Renshaw SA, Vollmer W, Foster SJ. *Staphylococcus aureus* cell wall structure and dynamics during host-pathogen interaction. PLoS Pathog. 17(3):e1009468. 2021.

37. Ledger EVK, Mesnage S, Edwards AM. Human serum triggers antibiotic tolerance in *Staphylococcus aureus*. Nat Commun. 13(1):2041. 2022.

38. Ellington JK, Harris M, Hudson MC, Vishin S, Webb LX, Sherertz R. Intracellular Staphylococcus aureus and antibiotic resistance: implications for treatment of staphylococcal osteomyelitis. J Orthop Res. 24(1):87–93. 2006.

39. Raineri EJM, Yedavally H, Salvati A, van Dijl JM. Time-resolved analysis of *Staphylococcus aureus* invading the endothelial barrier. Virulence. 11(1):1623–1639. 2020.

40. Reed P, Atilano ML, Alves R, Hoiczyk E, Sher X, Reichmann NT, Pereira PM, Roemer T, Filipe SR, Pereira-Leal JB, Ligoxygakis P, Pinho MG. *Staphylococcus aureus* Survives with a Minimal Peptidoglycan Synthesis Machine but Sacrifices Virulence and Antibiotic Resistance. PLoS Pathog. 11(5):e1004891. 2015.

41. Hines KM, Alvarado G, Chen X, Gatto C, Pokorny A, Alonzo F 3rd, Wilkinson BJ, Xu L. Lipidomic and Ultrastructural Characterization of the Cell Envelope of *Staphylococcus aureus* Grown in the Presence of Human Serum. mSphere. 5(3):e00339-20. 2020.

42. Yarwood JM, McCormick JK, Paustian ML, Kapur V, Schlievert PM. Repression of the *Staphylococcus aureus* accessory gene regulator in serum and *in vivo* . J Bacteriol . 184(4):1095–101. 2002.

43. James EH, Edwards AM, Wigneshweraraj S. Transcriptional downregulation of agr expression in *Staphylococcus aureus* during growth in human serum can be overcome by constitutively active mutant forms of the sensor kinase AgrC. FEMS Microbiol Lett . 349(2):153–62. 2013.

44. Manifold-Wheeler BC, Elmore BO, Triplett KD, Castleman MJ, Otto M, Hall PR. Serum Lipoproteins Are Critical for Pulmonary Innate Defense against *Staphylococcus aureus* Quorum Sensing. J Immunol. 196(1):328–35. 2016.

45. Peterson MM, Mack JL, Hall PR, Alsup AA, Alexander SM, Sully EK, Sawires YS, Cheung AL, Otto M, Gresham HD. Apolipoprotein B Is an innate barrier against invasive *Staphylococcus aureus* infection. Cell Host Microbe. 4(6):555–66. 2008.

46. Hall PR, Elmore BO, Spang CH, Alexander SM, Manifold-Wheeler BC, Castleman MJ, Daly SM, Peterson MM, Sully EK, Femling JK, Otto M, Horswill AR, Timmins GS, Gresham HD. Nox2 modification of LDL is essential for optimal apolipoprotein B-mediated control of agr type III *Staphylococcus aureus* quorum-sensing. PLoS Pathog. 9(2):e1003166. 2013.

47. Malachowa N, Whitney AR, Kobayashi SD, Sturdevant DE, Kennedy AD, Braughton KR, Shabb DW, Diep BA, Chambers HF, Otto M, DeLeo FR. Global changes in *Staphylococcus aureus* gene expression in human blood. PLoS One. 6(4):e18617. 2011.

48. Cybulska J, Jeljaszewicz J. Bacteriostatic activity of serum against staphylococci. J Bacteriol. 91(3):953–62. 1966.

49. Ehrenkranz NJ, Elliott DF, Zarco R. Serum Bacteriostasis of *Staphylococcus aureus* . Infect Immun. 3(5):664–70. 1971.

50. Painter KL, Hall A, Ha KP, Edwards AM. The Electron Transport Chain Sensitizes *Staphylococcus aureus* and *Enterococcus faecalis* to the Oxidative Burst. Infect Immun . 85(12):e00659–17. 2017.

51. Ha KP, Clarke RS, Kim GL, Brittan JL, Rowley JE, Mavridou DAI, Parker D, Clarke TB, Nobbs AH, Edwards AM. Staphylococcal DNA Repair Is Required for Infection. mBio. 11(6):e02288–20. 2020.

52. Fey PD, Endres JL, Yajjala VK, Widhelm TJ, Boissy RJ, Bose JL, Bayles KW. A genetic resource for rapid and comprehensive phenotype screening of nonessential *Staphylococcus aureus* genes. mBio. 4(1):e00537–12. 2013.

53. Horsburgh MJ, Aish JL, White IJ, Shaw L, Lithgow JK, Foster SJ. sigmaB modulates virulence determinant expression and stress resistance: characterization of a functional *rsbU* strain derived from *Staphylococcus aureus* 8325-4. J Bacteriol. 184(19):5457–67. 2002.

54. Gill SR, Fouts DE, Archer GL, Mongodin EF, Deboy RT, Ravel J, Paulsen IT, Kolonay JF, Brinkac L, Beanan M, Dodson RJ, Daugherty SC, Madupu R, Angiuoli SV, Durkin AS, Haft DH, Vamathevan J, Khouri H, Utterback T, Lee C, Dimitrov G, Jiang L, Qin H, Weidman J, Tran K, Kang K, Hance IR, Nelson KE, Fraser CM. Insights on evolution of virulence and resistance from the complete genome analysis of an early methicillin-resistant *Staphylococcus aureus* strain and a biofilm-producing methicillin-resistant *Staphylococcus epidermidis* strain. J Bacteriol. 2005 187(7):2426–38. 2005.

55. Baba T, Bae T, Schneewind O, Takeuchi F, Hiramatsu K. Genome sequence of *Staphylococcus aureus* strain Newman and comparative analysis of staphylococcal genomes: polymorphism and evolution of two major pathogenicity islands. J Bacteriol . 190(1):300–10. 2008.

56. Painter KL, Hall A, Ha KP, Edwards AM. The Electron Transport Chain Sensitizes *Staphylococcus aureus* and *Enterococcus faecalis* to the Oxidative Burst. Infect Immun . 85(12):e00659–17. 2017.

57. Nagl M, Kacani L, Müllauer B, Lemberger EM, Stoiber H, Sprinzl GM, Schennach H, Dierich MP. Phagocytosis and killing of bacteria by professional phagocytes and dendritic cells. Clin Diagn Lab Immunol. 9(6):1165–8. 2002.

58. Wonfor T, Li S, Dunphy RW, Macpherson A, van den Elsen J, Laabei M. Novel method for detecting complement C3 deposition on *Staphylococcus aureus*. Sci Rep. 12(1):15766. 2022.

59. Boero E, Brinkman I, Juliet T, van Yperen E, van Strijp JAG, Rooijakkers SHM, van Kessel KPM. Use of Flow Cytometry to Evaluate Phagocytosis of *Staphylococcus aureus* by Human Neutrophils. Front Immunol. 12:635825. 2021.

60. Silver LL. Fosfomycin: Mechanism and Resistance. Cold Spring Harb Perspect Med. 7:a025262. 2017.

61. Pujol M, Miró JM, Shaw E, Aguado JM, San-Juan R, Puig-Asensio M, Pigrau C, Calbo E, Montejo M, Rodriguez-Álvarez R, Garcia-Pais MJ, Pintado V, Escudero-Sánchez R, Lopez-Contreras J, Morata L, Montero M, Andrés M, Pasquau J, Arenas MD, Padilla B, Murillas J, Jover-Sáenz A, López-Cortes LE, García-Pardo G, Gasch O, Videla S, Hereu P, Tebé C, Pallarès N, Sanllorente M, Domínguez MÁ, Càmara J, Ferrer A, Padullés A, Cuervo G, Carratalà J; MRSA Bacteremia (BACSARM) Trial Investigators. Daptomycin Plus Fosfomycin Versus Daptomycin Alone for Methicillin-resistant *Staphylococcus aureus* Bacteremia and Endocarditis: A Randomized Clinical Trial. Clin Infect Dis. 72(9):1517–1525. 2021.

62. Wright AJ, Wilkowske CJ. The penicillins. Mayo Clin Proc. 66(10):1047–63. 1991.

63. Karlowsky JA, Kaplan N, Hafkin B, Hoban DJ, Zhanel GG. AFN-1252, a FabI inhibitor, demonstrates a *Staphylococcus*-specific spectrum of activity. Antimicrob Agents Chemother . 53(8):3544–8. 2009.

64. Sanders CC. Ciprofloxacin: in vitro activity, mechanism of action, and resistance. Rev Infect Dis. 10(3):516–27. 1988.

65. Douglas EJA, Palk N, Brignoli T, Altwiley D, Boura M, Laabei M, Recker M, Cheung GCY, Liu R, Hseih RC, Otto M, O’Brien E, McLoughlin RM, Massey RC. Extensive remodelling of the cell wall during the development of *Staphylococcus aureus* bacteraemia. Elife. 12:RP87026. 2023.

66. Rajagopal M, Walker S. Envelope Structures of Gram-Positive Bacteria. Curr Top Microbiol Immunol. 404:1–44. 2017.

67. Gautam S, Kim T, Lester E, Deep D, Spiegel DA. Wall teichoic acids prevent antibody binding to epitopes within the cell wall of *Staphylococcus aureus*. ACS Chem Biol. 11(1):25–30. 2016.

68. Leonard AC, Goncheva MI, Gilbert SE, Shareefdeen H, Petrie LE, Thompson LK, Khursigara CM, Heinrichs DE, Cox G. Autolysin-mediated peptidoglycan hydrolysis is required for the surface display of *Staphylococcus aureus* cell wall-anchored proteins. Proc Natl Acad Sci U S A. 120(12):e2301414120. 2023.

69. Miller LS, Fowler VG, Shukla SK, Rose WE, Proctor RA. Development of a vaccine against *Staphylococcus aureus* invasive infections: Evidence based on human immunity, genetics and bacterial evasion mechanisms. FEMS Microbiol Rev. 44(1):123–153. 2020.

70. Tsai CM, Caldera JR, Hajam IA, Chiang AWT, Tsai CH, Li H, Díez ML, Gonzalez C, Trieu D, Martins GA, Underhill DM, Arditi M, Lewis NE, Liu GY. Non-protective immune imprint underlies failure of *Staphylococcus aureus* IsdB vaccine. Cell Host Microbe . 30(8):1163–1172.e6. 2022.

71. Falcón R, Martínez A, Albert E, Madrid S, Oltra R, Giménez E, Soriano M, Vinuesa V, Gozalbo D, Gil ML, Navarro D. High vancomycin MICs within the susceptible range in *Staphylococcus aureus* bacteraemia isolates are associated with increased cell wall thickness and reduced intracellular killing by human phagocytes. Int J Antimicrob Agents. 47(5):343–50. 2016.

72. Tickle ARH, Ledger EVK, Edwards AM. Human serum induces daptomycin tolerance in *Enterococcus faecalis* and viridans group streptococci. Microbiology. 168(12). 2022.

73. Malachowa N, Whitney AR, Kobayashi SD, Sturdevant DE, Kennedy AD, Braughton KR, Shabb DW, Diep BA, Chambers HF, Otto M, DeLeo FR. Global changes in *Staphylococcus aureus* gene expression in human blood. PLoS One. 6(4):e18617. 2011.

74. Kim HK, Missiakas D, Schneewind O. Mouse models for infectious diseases caused by *Staphylococcus aureus*. J Immunol Methods. 410:88–99. 2014.

75. Foster TJ. Surface Proteins of *Staphylococcus aureus*. Microbiol Spectr. 7(4). 2019.

76. Luna BM, Nielsen TB, Cheng B, Pantapalangkoor P, Yan J, Boyle-Vavra S, Bruhn KW, Montgomery C, Spellberg B, Daum R. Vaccines targeting *Staphylococcus aureus* skin and bloodstream infections require different composition. PLoS One. 14(6):e0217439. 2019.

77. Clegg J, Soldaini E, McLoughlin RM, Rittenhouse S, Bagnoli F, Phogat S. *Staphylococcus aureus* Vaccine Research and Development: The Past, Present and Future, Including Novel Therapeutic Strategies. Front Immunol. 12:705360. 2021.

78. Marrie TJ, Costerton JW. Scanning and transmission electron microscopy of in situ bacterial colonization of intravenous and intraarterial catheters. J Clin Microbiol. 19(5):687–93. 1984.

79. Hetem DJ, de Ruiter SC, Buiting AGM, Kluytmans JAJW, Thijsen SF, Vlaminckx BJM, Wintermans RGF, Bonten MJM, Ekkelenkamp MB. Preventing *Staphylococcus aureus* bacteremia and sepsis in patients with Staphylococcus aureus colonization of intravascular catheters: a retrospective multicenter study and meta-analysis. Medicine. 90(4):284–288. 2011.

80. Minejima E, Mai N, Bui N, Mert M, Mack WJ, She RC, Nieberg P, Spellberg B, Wong-Beringer A. Defining the Breakpoint Duration of *Staphylococcus aureus* Bacteremia Predictive of Poor Outcomes. Clin Infect Dis. 70(4):566–573. 2020.

81. Ong PY, Ohtake T, Brandt C, Strickland I, Boguniewicz M, Ganz T, Gallo RL, Leung DY. Endogenous antimicrobial peptides and skin infections in atopic dermatitis. N Engl J Med . 347(15):1151–60. 2002.

82. Nizet V, Ohtake T, Lauth X, Trowbridge J, Rudisill J, Dorschner RA, Pestonjamasp V, Piraino J, Huttner K, Gallo RL . Innate antimicrobial peptide protects the skin from invasive bacterial infection. Nature. 414(6862):454–7. 2001.

83. Dorschner RA, Pestonjamasp VK, Tamakuwala S, Ohtake T, Rudisill J, Nizet V, Agerberth B, Gudmundsson GH, Gallo RL. Cutaneous injury induces the release of cathelicidin antimicrobial peptides active against group A *Streptococcus*. J. Invest. Dermatol. 117, 91–97 (2001).

84. Burian M, Plange J, Schmitt L, Kaschke A, Marquardt Y, Huth L, Baron JM, Hornef MW, Wolz C, Yazdi AS. Adaptation of *Staphylococcus aureus* to the Human Skin Environment Identified Using an *ex vivo* Tissue Model. Front Microbiol. 12:728989. 2021.

85. Kraus D, Herbert S, Kristian SA, Khosravi A, Nizet V, Götz F, Peschel A. The GraRS regulatory system controls *Staphylococcus aureus* susceptibility to antimicrobial host defenses. BMC Microbiol. 8:85. 2008.

86. Cheung AL, Cho J, Bayer AS, Yeaman MR, Xiong YQ, Donegan NP, Mikheyeva IV, Lee GY, Yang SJ. Role of the *Staphylococcus aureus* Extracellular Loop of GraS in Resistance to Distinct Human Defense Peptides in PMN and Invasive Cardiovascular infections. Infect Immun. 89(10):e0034721. 2021.

87. Friberg C, Haaber JK, Vestergaard M, Fait A, Perrot V, Levin BR, Ingmer H. Human antimicrobial peptide, LL-37, induces non-inheritable reduced susceptibility to vancomycin in *Staphylococcus aureus*. Sci Rep. 10(1):13121. 2020.

88. Ranganathan N, Johnson R, Edwards AM. The general stress response of *Staphylococcus aureus* promotes tolerance of antibiotics and survival in whole human blood. Microbiology. 166(11):1088–1094. 2020.

89. Rowe SE, Wagner NJ, Li L, Beam JE, Wilkinson AD, Radlinski LC, Zhang Q, Miao EA, Conlon BP. Reactive oxygen species induce antibiotic tolerance during systemic *Staphylococcus aureus* infection. Nat Microbiol. 5(2):282–290. 2020.

90. Beam JE, Rowe SE, Conlon BP. Shooting yourself in the foot: How immune cells induce antibiotic tolerance in microbial pathogens. PLoS Pathog. 17(7):e1009660. 2021.

91. Shen F, Tang X, Cheng W, Wang Y, Wang C, Shi X, An Y, Zhang Q, Liu M, Liu B, Yu L. Fosfomycin enhances phagocyte-mediated killing of *Staphylococcus aureus* by extracellular traps and reactive oxygen species. Sci Rep. 6:19262. 2016.

92. Berti A, Rose W, Nizet V, Sakoulas G. Antibiotics and Innate Immunity: A Cooperative Effort Toward the Successful Treatment of Infections. Open Forum Infect Dis. 7(8):ofaa302. 2020.

93. Fey PD, Endres JL, Yajjala VK, Widhelm TJ, Boissy RJ, Bose JL, Bayles KW. A genetic resource for rapid and comprehensive phenotype screening of nonessential *Staphylococcus aureus* genes. mBio. 4(1):e00537–12. 2013.

94. Higgins J., Loughman A., van Kessel K. P., van Strijp J. A., Foster T. J. Clumping factor A of *Staphylococcus aureus* inhibits phagocytosis by human polymorphonuclear leucocytes. FEMS Microbiol. Lett. 258:290–296. 2006.

95. Olsen CH. Review of the use of statistics in infection and immunity. Infect Immun. 1(12):6689–92. 2003.

96. Cunnion KM, Hair PS, Buescher ES. Cleavage of complement C3b to iC3b on the surface of *Staphylococcus aureus* is mediated by serum complement factor I. Infect Immun. 72(5):2858–63. 2004.

97. Easmon CS, Lanyon H, Cole PJ. Use of lysostaphin to remove cell-adherent staphylococci during *in vitro* assays of phagocyte function. Br J Exp Pathol. 59(4):381–5. 1978.

98. Simon GL, Miller HG, Borenstein DG. Synovial fluid inhibits killing of *Staphylococcus aureus* by neutrophils. Infect Immun. 40(3):1004–10. 1983.

99. Schneewind O, Mihaylova-Petkov D, Model P. Cell wall sorting signals in surface proteins of gram-positive bacteria. EMBO J. 12(12):4803–11. 1993.

