## Supplementary figures for "Host-induced cell wall remodelling impairs opsonophagocytosis of *Staphylococcus aureus* by neutrophils"

**Staphylococcal host adaptation impairs opsonisation by antibody and complement**

**Supplementary data file**

Supplementary Figures S1-S9.

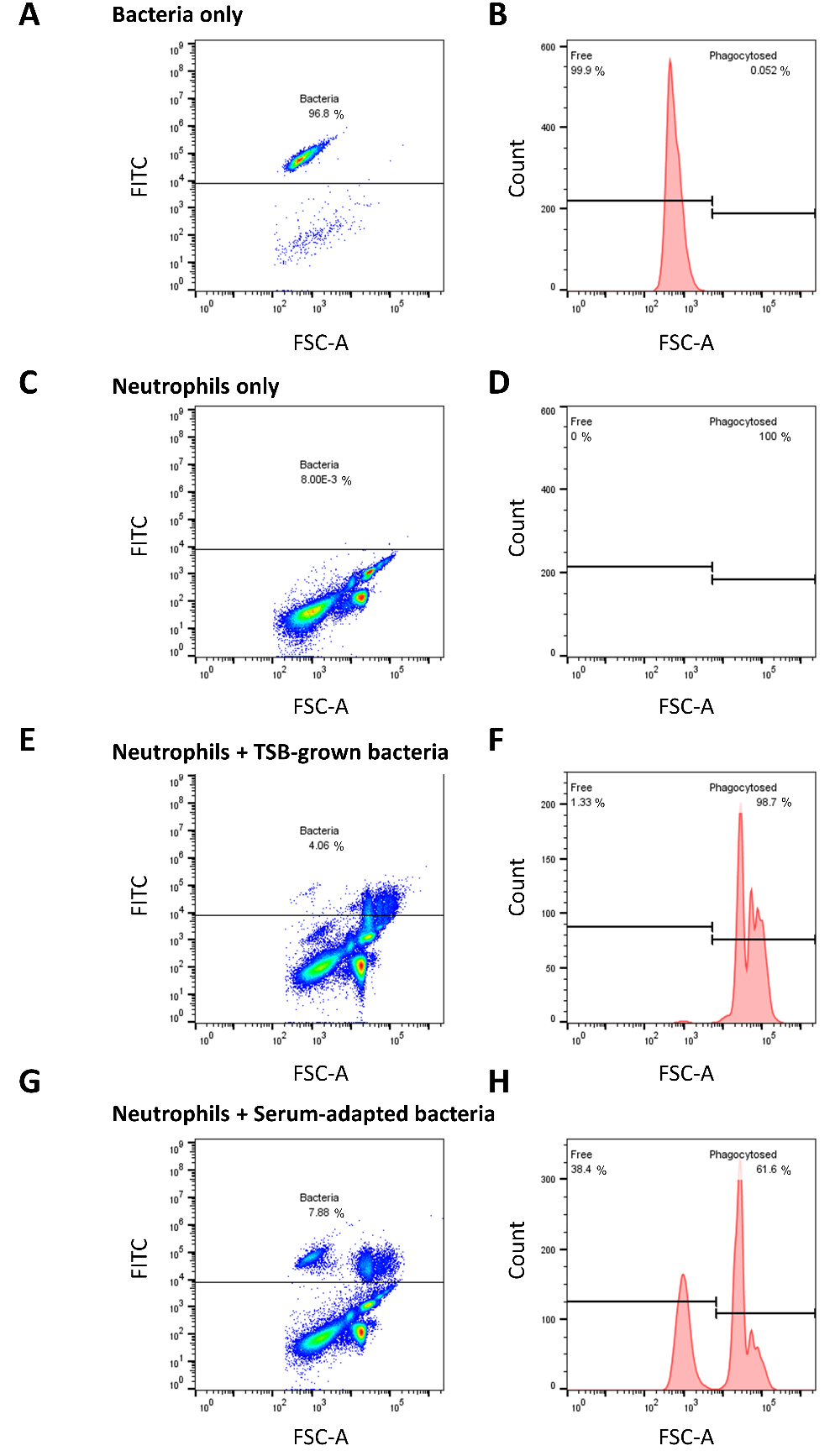

**Supplementary figure S1. Gating strategy for flow cytometry-based opsonophagocytosis assay.** Bacteria were labelled with FITC and gated based on fluorescence (A) and FSC (B). Neutrophils were not fluorescent (C) and had a greater FSC than bacteria (D). This enabled us to detect a shift in FSC of fluorescent bacteria when they were phagocytosed, which happened at high efficiency for broth grown bacteria (E, F). By contrast, many host-adapted bacteria remained un-phagocytosed by neutrophils (G, H).

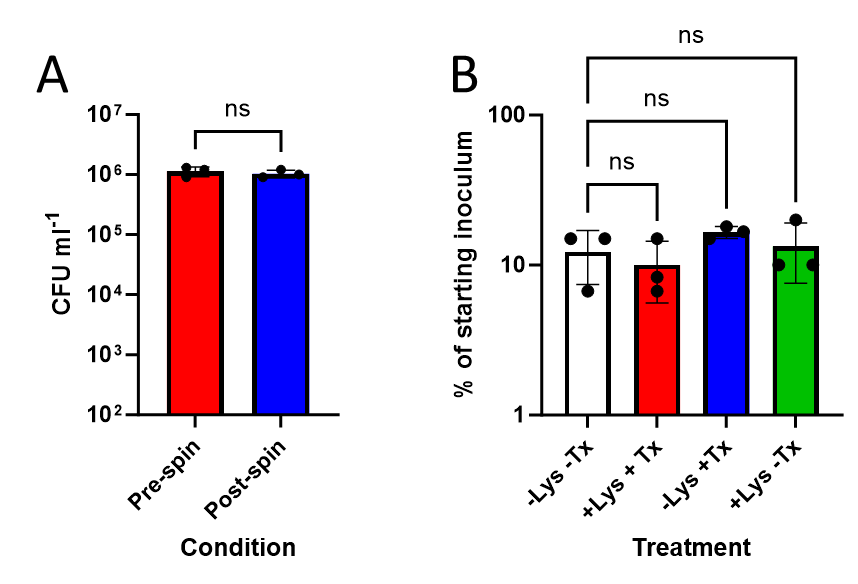

**Supplementary figure S2. Neutrophil-associated bacteria are intracellular.** One of the phagocytosis assays used relies on a centrifugation step (500 x *g*, 1 min) to separate free bacteria from those associated with neutrophils. To demonstrate that this centrifugation step was appropriate, *S. aureus* cells were suspended in HBSS and then centrifuged at 500 x *g* (1 min). CFU counts in the supernatant before and after centrifugation were determined for three independent experiments (**A**). This demonstrated that this centrifugation step does not pull-down free bacteria out of suspension.

Since bacteria that are pulled down via association with neutrophils could be phagocytosed or attached to the outside, we examined the impact of lysostaphin (Lys) which will kill extracellular but not intracellular bacteria [88,89], on CFU counts of bacteria associated with neutrophils (**B**). Since the presence of lysostaphin had no effect on CFU counts of bacteria associated with neutrophils, we can conclude that neutrophil-associated bacteria have been phagocytosed. We also determined whether lysis of neutrophils with Triton X-100 (Tx) increased recovery of phagocytosed bacteria (**B**). However, the presence of the detergent had no effect on CFU counts of bacteria phagocytosed by neutrophils.

In (**A**), bars show the mean of 3 independent experiments. Error bars show the standard deviation. Data were analysed by student’s t-test. In (**B**) bars show the mean of 3 independent experiments. Error bars show the standard deviation. Data were analysed by one-way ANOVA and Dunnett’s post hoc test for multiple comparisons (ns p>0.05).

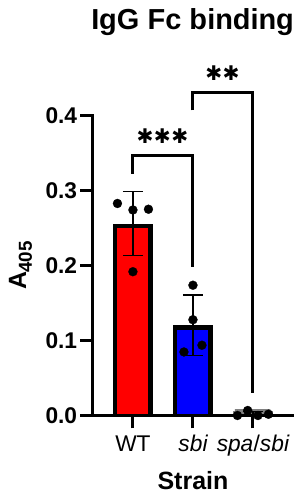

**Supplementary figure S3. A Δ*spa* *sbi*::Tn double mutant does not bind IgG Fc.** To ensure that the constructed *S. aureus* JE2 Δ*spa* *sbi*::Tn double mutant did not bind IgG, we measured the adhesion of biotinylated IgG Fc fragment to the surface of *S. aureus* JE2 wild type (WT) and mutant strains. As expected, wild type *S. aureus* bound high levels of IgG Fc whilst a *sbi*::Tn mutant showed reduced binding and there was no detectable binding by the Δ*spa* *sbi*::Tn double mutant. Bars show the mean of 4 independent experiments. Error bars show the standard deviation. Data were analysed by one-way ANOVA and Dunnett’s post hoc test for multiple comparisons (**p=<0.01, ***p=<0.001).

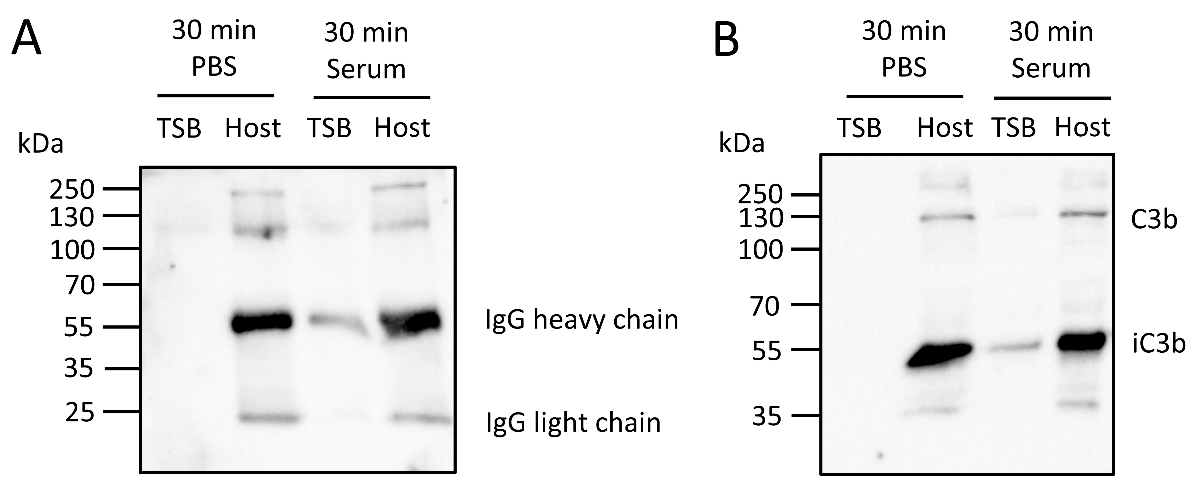

**Supplementary figure S4. Host-adapted bacteria have large amounts of bound antibody and complement****.** TSB-grown and serum-adapted *S. aureus* cells were incubated for 30 min with PBS or 10 % fresh serum before the levels of bound (**A**) IgG and (**B**) C3 were determined by western blotting. The expected positions for IgG heavy and light chains (**A**) and C3b and ic3b (**B**) are indicated. As reported previously, we observed processing of C3b to iC3b on the *S. aureus* cell surface [64].

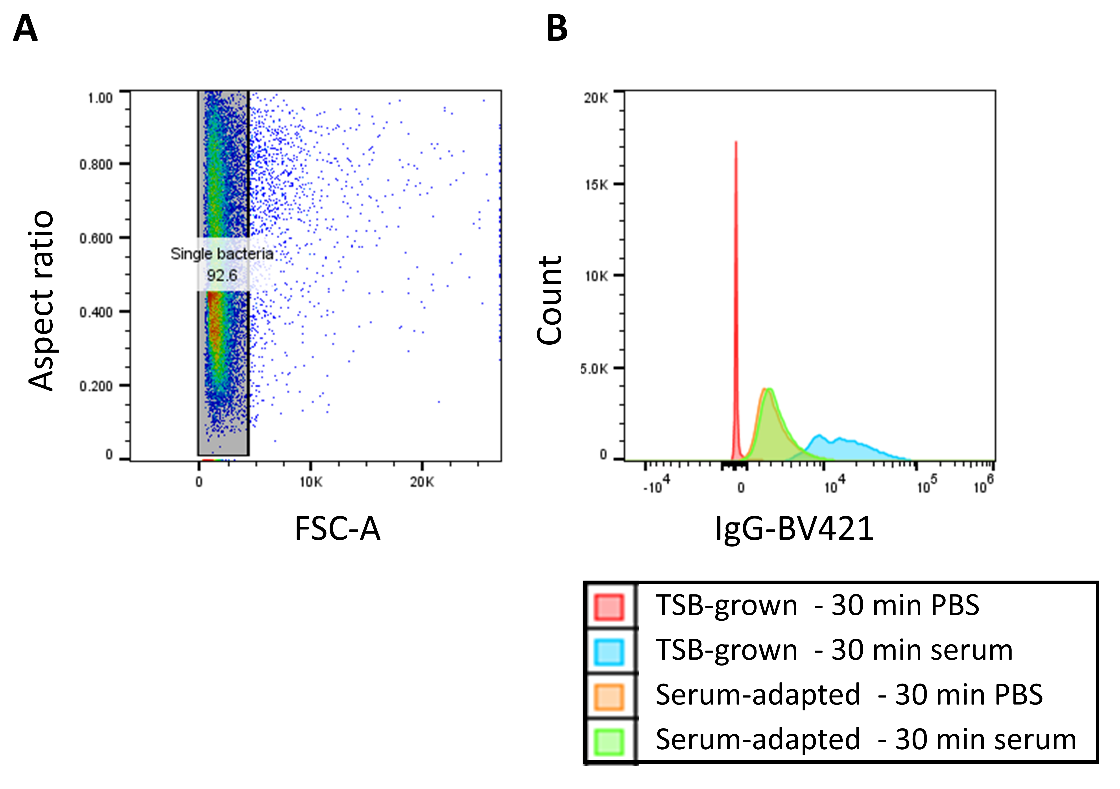

**Supplementary figure S5. Gating strategy for flow cytometry-based determination of opsonin exposure.** Bacteria were grown in broth or allowed to adapt to human serum before labelling with antibiotics against complement C3 component or human antibody. Bacteria were gated based on FSC values to select single cells (A). The fluorescence of each single cell was then measured to generate median fluorescence values for each experimental condition. For example, bacteria grown in broth, or adapted to host conditions by incubation in serum and then exposed, or not, to fresh serum (B). This shows that, for broth-grown bacteria that had not been exposed to serum there was no fluorescence, indicative of an absence of non-specific binding. However, for broth grown bacteria exposed to serum there was a strong fluorescence signal (B). Host adapted bacteria were moderately fluorescent, regardless of whether they had been exposed to fresh serum or not (B).

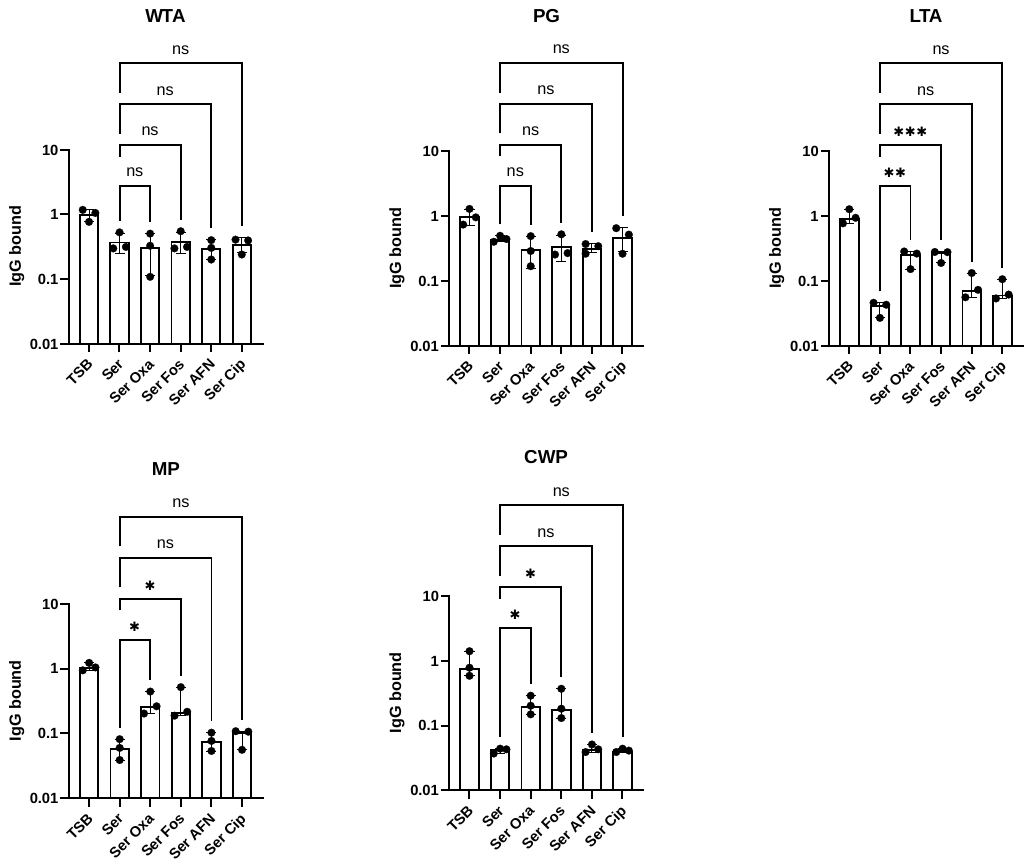

**Supplementary figure S6. Antibiotics that inhibit cell wall synthesis promote accessibility of bound IgG against LTA and surface proteins during host adaptation.** *S. aureus* was grown in TSB to exponential phase before incubation in serum for 30 min to enable opsonisation to occur (TSB), or for 16 h to allow host adaptation (Ser) +/- the following antibiotics: oxacillin (Oxa), fosfomycin (Fos), AFN-1252 (AFN) or ciprofloxacin (Cip). Subsequently, surface exposed IgG was eluted and then assayed for binding to major surface structures wall teichoic acid (WTA), peptidoglycan (PG), lipoteichoic acid (LTA), membrane-associated proteins (MP) or cell wall-associated proteins (CWP). Data were normalised to values for the TSB condition and represent the median ± 95% CI of three independent biological replicates and were analysed by Kruskal Wallis test and Dunn’s *post-hoc* test to establish statistically significant differences between groups (***, P <0.001; **, P < 0.01; *, P < 0.05; ns, P ≥ 0.05 for the indicated comparisons).

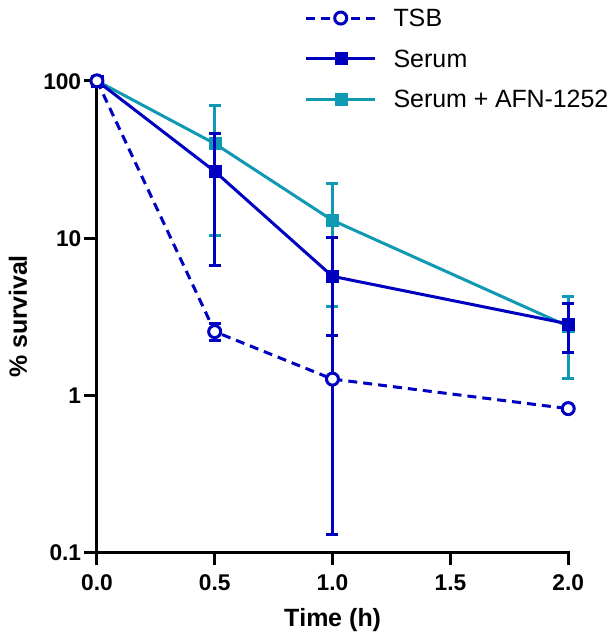

**Supplementary figure S7. Host adaptation in the presence of AFN-1252 does not lead to increased phagocytic killing of *S. aureus*.** Survival of TSB-grown (TSB) and serum-adapted cultures of wild type *S. aureus* USA300 (Serum) or serum-adapted in the presence of AFN-1252, which inhibits fatty acid synthesis. Data represent the mean ± standard deviation of three independent biological replicates. Data were analysed by a two-way ANOVA with Sidak’s *post-hoc* test (the presence of AFN-1252 during host adaptation did not affect susceptibility to neutrophil-mediated killing P > 0.05).

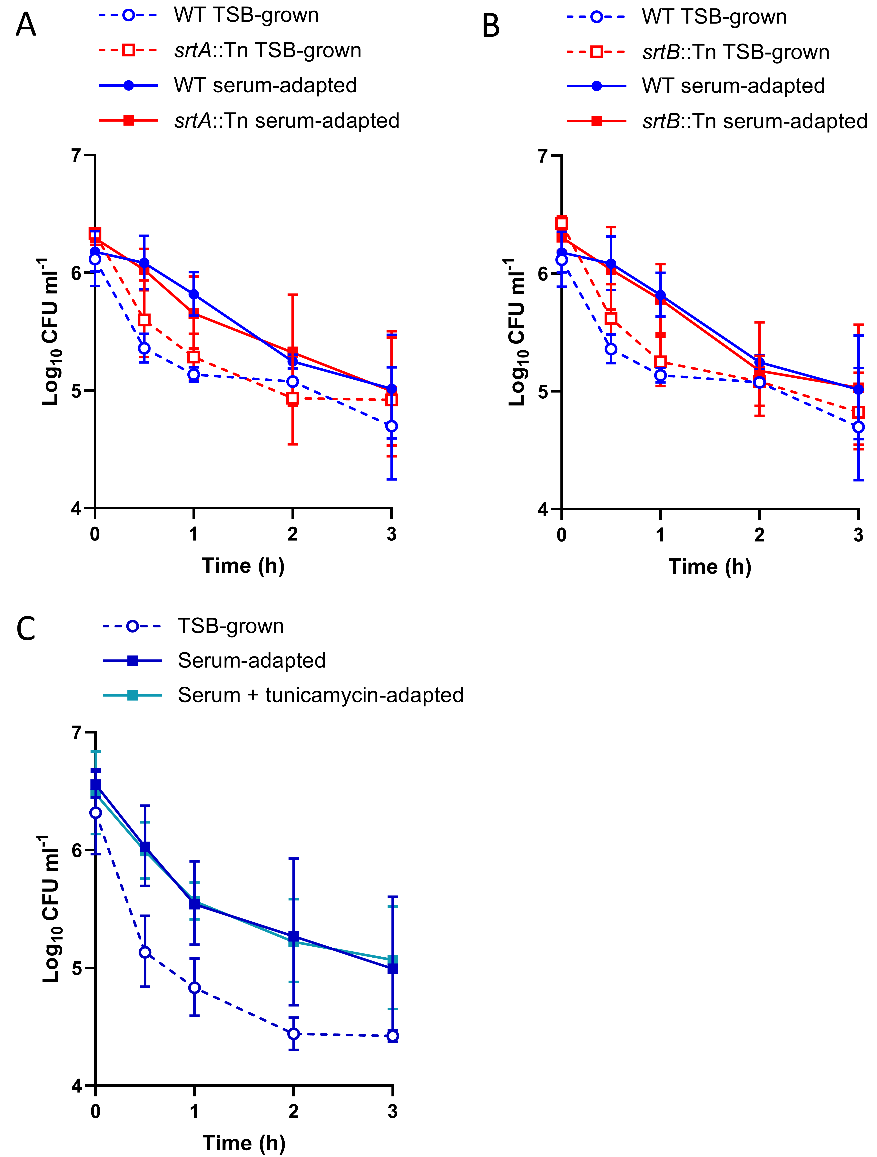

**Supplementary figure S8. Peptidoglycan-anchored proteins do not contribute to immune evasion in serum-adapted *S. aureus*.** Wild type and mutant bacteria were grown in broth or allowed to adapt to human serum before incubation with human neutrophils in the presence of serum and staphylococcal survival measured over time. (**A**), Log10 CFU ml^-1^ survival of TSB-grown and serum-adapted cultures of wild type *S. aureus* USA300 or a *srtA*::Tn mutant that lacks peptidoglycan-bound LPXTG-motif proteins. (**B**), Log10 CFU ml^-1^ survival of TSB-grown and serum-adapted cultures of wild type *S. aureus* USA300 or a *srtB*::Tn mutant that lacks peptidoglycan-bound NPQTN-motif protein IsdC. Data were analysed by a two-way ANOVA with Sidak’s *post-hoc* test (neither mutant was significantly affected for survival compared to wild type, P = > 0.05). All data were generated in the same experiments, but values for *srtA*::Tn and *srtB*::Tn are shown on separate graphs for clarity. Data for WT cells are the same in both graphs.

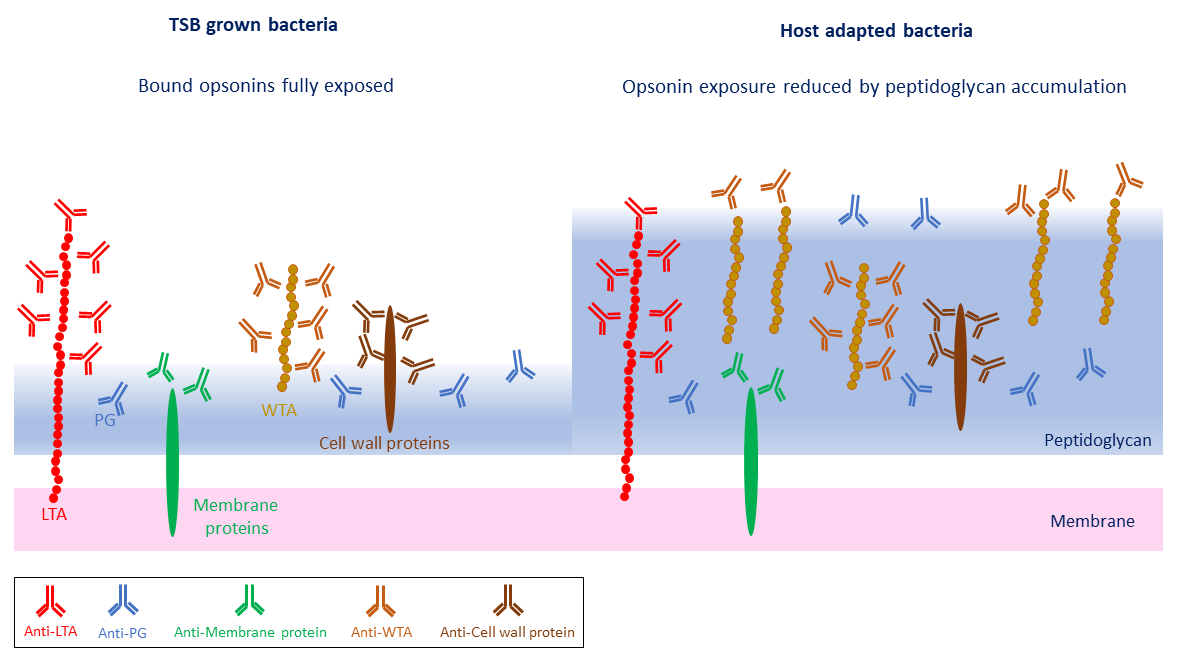

**Supplementary figure S9. Summary diagram of how the adaptation of *S. aureus* to the host environment modulates the exposure of bound opsonins.** TSB grown bacteria have high levels of exposed surface structures and antibodies that target them, resulting in efficient opsonophagocytic killing. However, as *S. aureus* adapts to the host, peptidoglycan accumulates, which conceals many of these bound antibodies, resulting in reduced opsonophagocytic killing.
